# The repeat region of the circumsporozoite protein is an elastic linear spring with a functional role in *Plasmodium* sporozoite motility

**DOI:** 10.1101/2021.05.12.443759

**Authors:** Amanda E. Balaban, Sachie Kanatani, Jaba Mitra, Jason Gregory, Natasha Vartak, Ariadne Sinnis-Bourozikas, Fredrich Frischknecht, Taekjip Ha, Photini Sinnis

**Affiliations:** Johns Hopkins Malaria Research Institute and Department of Molecular Microbiology & Immunology, Johns Hopkins Bloomberg School of Public Health, Baltimore, MD 21201; Department of Biophysics and Biophysical Chemistry, Johns Hopkins University, Baltimore, MD 21218; Department of Materials Science and Engineering, University of Illinois at Urbana-Champaign, IL 61801; Integrative Parasitology, Center for Infectious Diseases, Heidelberg University Medical School, Heidelberg, Germany; German Center for Infection Research (DZIF), Heidelberg, Germany; Howard Hughes Medical Institute, Johns Hopkins University, Baltimore, MD

**Keywords:** malaria, circumsporozoite protein, motility, repeats, sporozoite

## Abstract

The circumsporozoite protein (CSP) forms a dense coat on the surface of the sporozoite, the infective stage of the malaria parasite. The central repeat region of CSP is a critical component of the only licensed malaria vaccine yet little is known about its structure or function. We found that sporozoite mutants with severely truncated or scrambled repeats have impaired motility due to altered adhesion site formation and dynamics, suggesting that the CSP repeats provide a cohesive environment in which adhesion sites can form. We hypothesized that biophysical properties of the repeats are important in this role and interrogated this using single-molecule fluorescence-force spectroscopy. We show that the repeats are a stiff, linear spring with elastic properties, dependent upon length and lost when the repeats are scrambled. These data are the first evidence that the CSP repeat region serves a functional role during infection and motility, likely mediated through its biophysical properties.

**Summary:** No clear function of the central repeat region of the malaria circumsporozoite protein has been described to date, despite its central role in the only licensed malaria vaccine. Here we use mutational analysis and single-molecule fluorescence-force spectroscopy to describe the structural properties and determine the function of this conserved region and important vaccine target.

**Highlights:** The CSP repeats have properties of a linear spring

Scrambling or large truncations of the repeats leads to defects in sporozoite motility

Motility defects are attributed to abnormal formation of adhesion sites

## Introduction

Every year there are 228 million new malaria cases resulting in an estimated 405,000 deaths, the majority of which are in children in Sub-Saharan Africa (WHO, 2019). Malaria is caused by protozoan parasites of the genus *Plasmodium*, whose complex life cycle requires cycling between mosquito and vertebrate hosts. Sporozoites, the infective stage of the parasite, develop in oocysts on the mosquito midgut wall and enter salivary glands, where they wait to be inoculated into the mammalian host as the infected mosquito probes for blood. Once inoculated, sporozoites migrate through the dermis to find and enter blood vessels. From here, they are carried to the liver and enter hepatocytes, where they develop and multiply, ultimately releasing thousands of hepatic merozoites and initiating the blood stage of infection. Iterative asexual replication cycles in the blood lead to the high parasite numbers responsible for all clinical symptoms of malaria. The pre-erythrocytic stages, sporozoites and liver stages, establish infection in the mammalian host and have proven to be a promising vaccine target, likely due to the bottleneck in parasite numbers during transmission.

Vaccines present the best long-term approach to malaria control and eventual eradication. The major surface protein of the sporozoite, the circumsporozoite protein (CSP), is the basis of the most advanced malaria vaccine to date, with RTS,S, a subunit vaccine based on CSP, demonstrating ∼48% protection from infection and severe disease for one year (The RTS,S Clinical Trials Partnership, 2011). Though falling short of community-established goals, with efficacy waning significantly after one year (Nussenzweig et al., 2011; Olotu et al., 2016), RTS,S validated the sporozoite and CSP as a vaccine target. RTS,S was developed based on early immunization studies in the rodent model, demonstrating that CSP is an immunodominant antigen (Cohen et al., 2010) and that protection correlated with antibodies to the CSP repeat region (Foquet et al., 2014). Thus, a better understanding of the function of CSP and its role in sporozoite biology could rationally guide the design of future vaccine candidates.

CSP forms a dense coat on the parasite’s surface and is composed of 3 domains (Figure 1A): An N-terminal domain ending in a conserved proteolytic cleavage site (Region I), a central repeat domain, and a carboxy-terminal adhesive domain, the type I thrombospondin repeat (TSR) (Coppi et al., 2011). Previous studies have shown that the N-terminal domain masks the TSR as sporozoites migrate from the mosquito midgut to the mammalian liver (Coppi et al., 2011). In the liver, a sporozoite protease cleaves CSP at Region I, leading to the removal of the N-terminus and exposure of the TSR, an event that is associated with the switch from a migratory to an invasive sporozoite (Coppi et al., 2011). Though these studies began to unravel the function of the N- and C-terminal domains of CSP, little functional work has been performed on the central repeat region.

**Figure 1:**
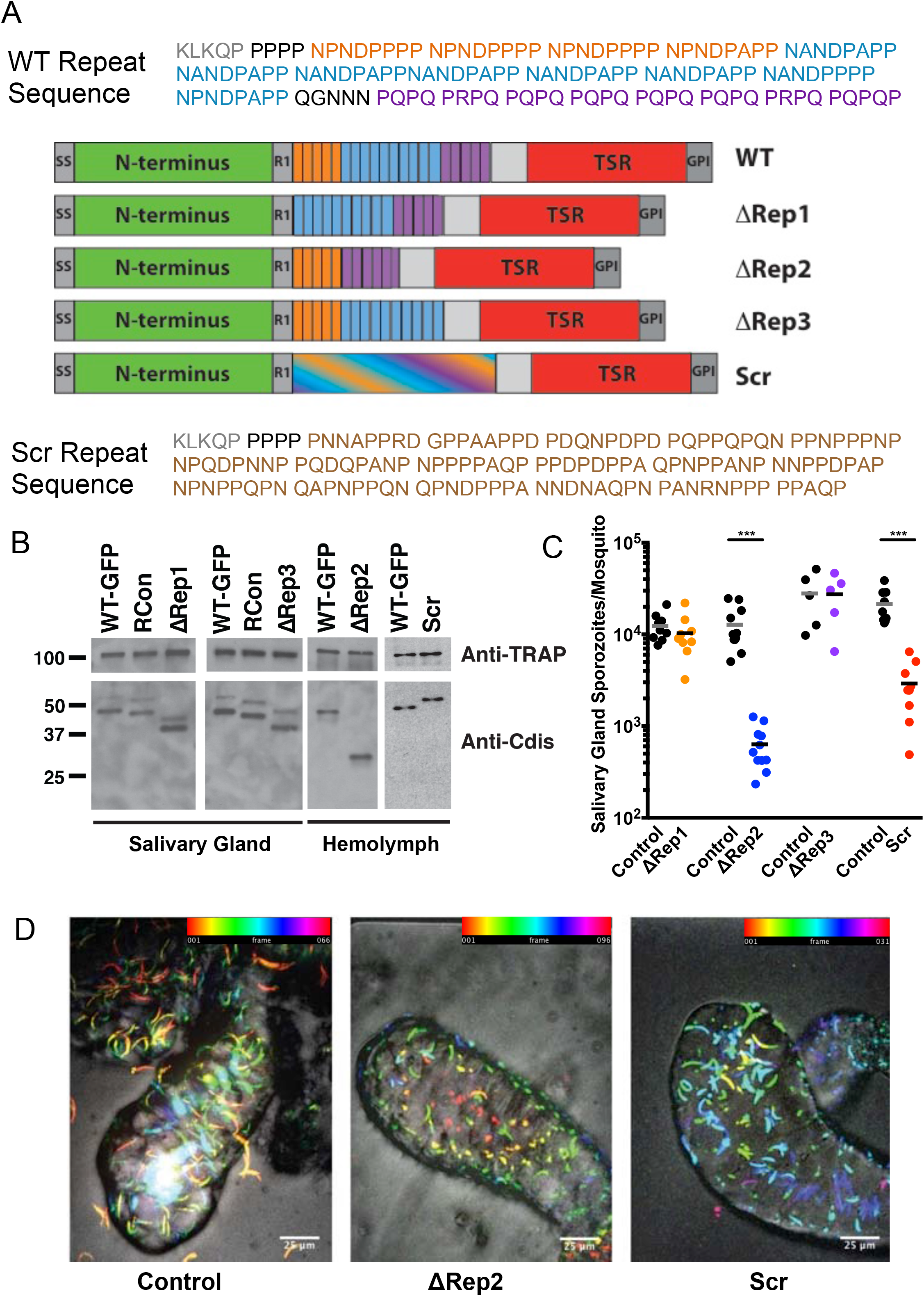
CSP Expression and Salivary Gland Invasion of Repeat Mutants. **(A)** Schematic of CSP repeat mutants. SS, signal sequence; R1, region I; TSR, type I thrombospondin repeat; GPI, putative glycosylphosphatidyl inositol anchor. The amino acid sequence above the diagram is the WT *Plasmodium berghei* repeat sequence, with the region I cleavage site in gray. The three repeated sequences are color coded: orange = first repeat, blue = second repeat, and purple = third repeat. The scrambled repeats are indicated with blending of the different repeat block colors and the scrambled sequence is shown below the diagram. **(B)** CSP expression in repeat mutants. Lysates of salivary gland or hemolymph repeat mutant and control sporozoites were subject to western blot analysis, using antisera specific for the disordered region in the C-terminus of CSP to visualize CSP expression, with TRAP antisera as a loading control. For ΔRep2 and Scr blots, hemolymph sporozoites were used because of the limited numbers of salivary gland sporozoites. Molecular weight markers are indicated on the left. **(C)** Salivary gland sporozoite invasion. Salivary glands from 20 infected mosquitoes were pooled, homogenized and the average number of sporozoites per mosquito was determined using a hemocytometer Each data point is from an independent mosquito cycle with matched WT-GFP and mutant cycles. Kruskal-Wallis p<0.0001. Mann-Whitney-Wilcoxon ***p<0.005. **(D)** Live confocal imaging of infected mosquito salivary glands of WT-GFP, ΔRep2, and Scr infected mosquitoes with GFP channel (sporozoites) and DIC. GFP channel is pseudo-colored with different colors representing different positions in Z, allowing for visualization of sporozoite distribution through depth of salivary gland tissue. Scale bars = 25 µm; Color bar scales on top right indicate section depth.

The repeat region of CSP consists of tandem amino acid repeats that vary among *Plasmodium* species (Supplementary Table 1) (Kemp et al., 1987). Interestingly, the repeats of different *Plasmodium* species are exchangeable, with no observable phenotype, as has been demonstrated with the generation of chimeric parasite lines (Espinosa et al., 2013; Persson et al., 2002). These data suggest that the repeats have a conserved structure and function across species despite variant sequences. Previous studies on the structure of the CSP repeats have used recombinant repeat peptides and provided evidence for a β-turn conformation (Plassmeyer et al., 2009; Ghasparian et al., 2006; Fasman et al., 1990; Dyson et al., 1990; Verdini et al., 1991). However, because these studies have not been conducted on full-length protein and no functional assays exist to confirm proper folding, the consensus in the field has been that the repeats are a disordered, non-functional region. To probe the function of CSP repeat region, we generated mutant parasites with truncated or altered repeats, in the rodent malaria model *Plasmodium berghei* and interrogated their biophysical properties using single-molecule fluorescence-force spectroscopy.

## Results

### Generation of CSP repeat region mutants

Previous work showed that deletion of the entire CSP repeat region results in severe defects in sporozoite development, with decreased oocyst sporozoite numbers and sporozoite death prior to exiting the oocyst (Ferguson et al., 2014). Since none of these sporozoites reached the salivary glands, it was not possible to elucidate the function of the repeats in the mammalian host using these lines. To overcome this limitation, we created mutants with more subtle phenotypes. We previously established that during migration within the mosquito, the exposure of the N- and C- terminal domains of CSP on the surface is altered to modify sporozoite adhesion (Coppi et al., 2011). To probe whether the length of the repeats was important for domain exposure, we generated three CSP repeat truncation mutants, ΔRep1, ΔRep2, and ΔRep3, in which different lengths or regions of the repeats were deleted (Figure 1A). Additionally, to test whether the repetitive nature of the repeats contains important structural features we also generated a scrambled mutant, Scr, by randomly scrambling the amino acid sequence of the repeats. For this mutant, we avoided introduction of new secondary structures, as predicted by the Chou-Fasman method, but maintained amino acid content and length (Figure 1A). These mutants were generated in *P. berghei* ANKA 507 clone 1, a line that constitutively expresses GFP and was selected by flow cytometry so that it does not contain a selection cassette (Janse et al., 2006). Transfection plasmids containing the desired *csp* repeat mutations were designed to replace the endogenous wildtype *csp* gene (Supplementary Figure 1). Transfected parasites were cloned and verified by a series of diagnostic PCRs and sequencing of the *csp* gene (Supplementary Figure 1). Phenotyping of CSP repeat mutants was always performed in parallel with a control: In some cases, this was the parental line *P. berghei* ANKA 507 clone 1 (WT-GFP) and in other cases this was a recombinant control line (RCon) in which a wildtype *csp* was transfected into the native locus, controlling for any effects that the altered genomic locus might have. Side by side comparisons of WT-GFP and RCon parasites showed that they had similar phenotypes in the assays used in our study (Supplementary Figure 2). Thus, throughout the manuscript, “Controls” refer to either WT-GFP or RCon parasites.

### ΔRep2 and Scr sporozoites do not progress normally through the mosquito

To characterize the progression of CSP repeat mutants through the mosquito stages, we first assessed the number of salivary gland sporozoites in each line. As shown in Figure 1C, ΔRep1 and ΔRep3 had comparable salivary gland infections to controls. In contrast, ΔRep2 and Scr parasites were inhibited in their capacity to enter the salivary glands, with fewer salivary gland associated sporozoites. We confirmed that ΔRep2 and Scr salivary gland associated sporozoites were inside the salivary glands, as opposed to attached to the external surface of the glands by live confocal imaging of salivary glands (Figure 1D).

Since ΔRep2 and Scr exhibited lower numbers of salivary gland sporozoites, we characterized their mosquito phenotypes in more detail. ΔRep2 and Scr sporozoite development in oocysts was unaltered compared to controls, with a comparable number of midgut sporozoites per mosquito in five independent mosquito cycles (Supplementary Figure 3). We then investigated whether ΔRep2 and Scr mutants had defects in egress from oocysts by quantifying sporozoite numbers in the hemolymph, the open circulatory system of the mosquito. We found that these mutants had similar numbers of hemolymph sporozoites as controls (Supplementary Figure 3). These data confirm that the decreased number of salivary gland sporozoites in ΔRep2 and Scr mutants is due to a salivary gland invasion defect and not an earlier developmental phenotype.

### Expression and conformation of mutant CSP

To determine whether CSP was expressed at similar levels in the repeat mutants as in control sporozoites, we performed western blot analysis with antisera generated to the disordered region just after the repeats. As shown in Figure 1B, mutant sporozoites expressed similar amounts of CSP, demonstrating that the repeat mutations did not alter CSP expression. To perform these experiments with the ΔRep2 and Scr mutants we used hemolymph sporozoites as the low number of salivary gland sporozoites resulted in large amounts of contaminating mosquito debris, which gave high background on western blots. We then went on to determine whether mutant CSP was conformationally altered, using previously developed polyclonal antisera specific for the C-terminal domain of CSP (Coppi et al., 2011). Previous studies found that the C-terminal adhesion domain was exposed only during sporozoite development in the oocysts and upon reaching the liver (Coppi et al., 2011). In between these two locations, the N-terminal domain masks the C-terminal domain (Coppi et al., 2011). By immunofluorescence microscopy, we found that salivary gland sporozoites of the ΔRep1, ΔRep3, and Scr CSP repeat mutants did not stain with the C-terminal antisera, similar to controls. In contrast, ΔRep2 sporozoites had enhanced C-terminal exposure (Figure 2A). Treatment of sporozoites with 0.1% saponin during staining with the anti-C-terminal antibody to break intramolecular interactions in the surface CSP and expose the C-terminal domain, allowed for the comparison of total CSP on the surface of mutant sporozoites. Using this method, we found comparable levels of CSP on the surface of control and mutant sporozoites (Figure 2B), confirming our western blot results.

**Figure 2:**
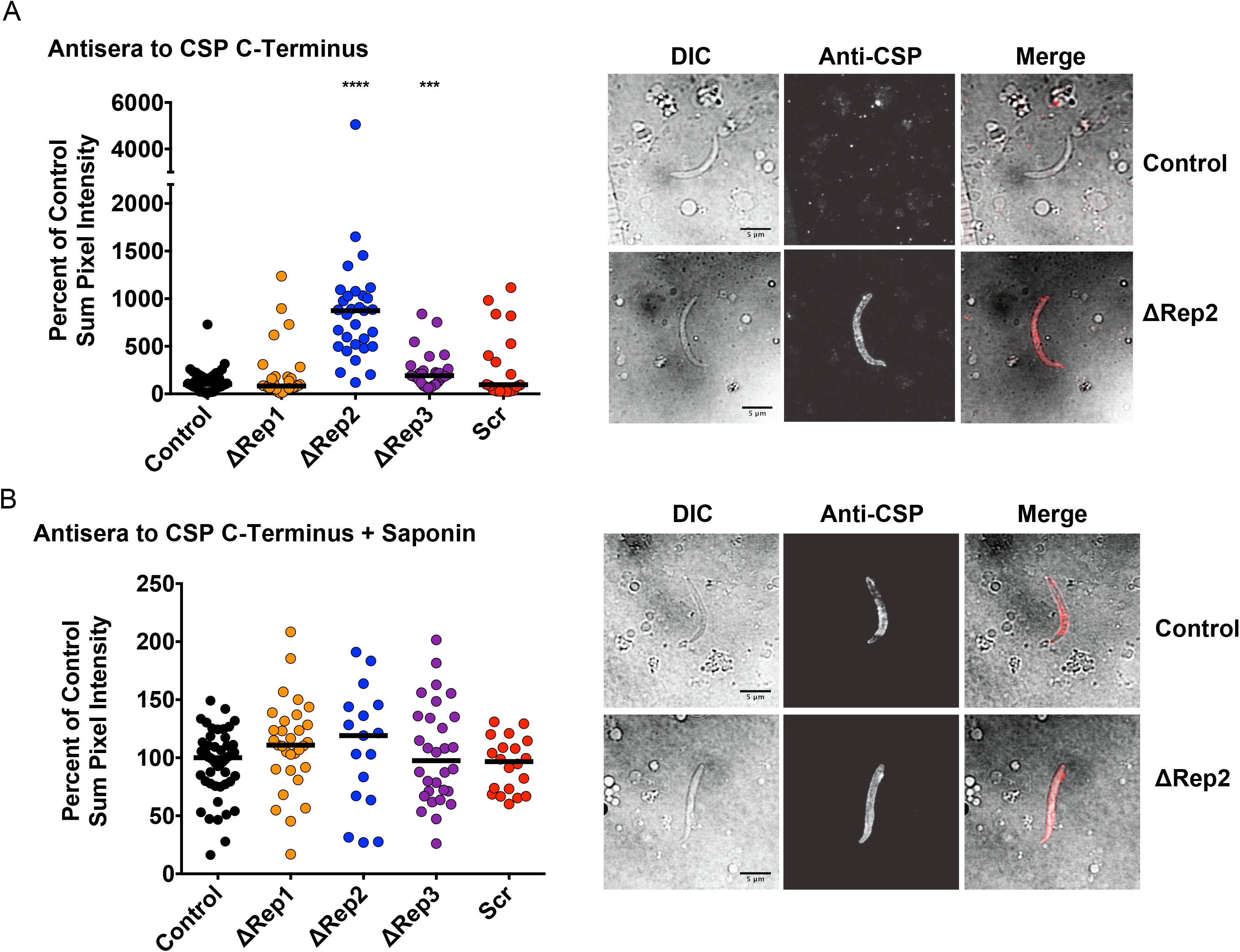
ΔRep2 sporozoites have enhanced exposure of the carboxy-terminal TSR. Salivary gland sporozoites of the indicated mutant lines were stained with antibodies specific to the C-terminus of CSP. Sporozoites were unpermeabilized **(A)** or, 0.1% saponin was included with the antisera **(B).** 20-30 sporozoites per condition were quantified for sum pixel intensity and presented as percent of the median pixel intensity of matched WT-GFP salivary gland sporozoites. For (A) Kruskal-Wallis p<0.0001 followed by Mann-Whitney- Wilcoxon ***p<0.0005, ****p<0.0001. For (B) Kruskal-Wallis p = 0.1980. Representative images of WT-GFP and ΔRep2 for each antibody condition are shown to the right of each panel with DIC, fluorescence and merged images. Scale bars = 5 µm.

The enhanced C-terminal exposure on ΔRep2 sporozoites could be explained by one of two mechanisms: 1) constitutive exposure of the CSP C-terminus on the surface of ΔRep2 sporozoites due to an inability of the N-terminus to mask the C-terminus in this mutant or 2) proteolytic processing of the N-terminus of CSP, which would lead to exposure of the C-terminal domain (Coppi et al., 2005, 2011). To determine if premature cleavage of the N-terminus could account for the exposure of the C-terminus in ΔRep2 sporozoites, we metabolically labeled CSP in control, ΔRep1 (as an additional control), and ΔRep2 salivary gland sporozoites to observe CSP cleavage. No differences in CSP cleavage were observed in ΔRep2 sporozoites (Supplementary Figure 4). Thus, we hypothesize that the repeats form a linker between the N- and C-terminal domains and must be of a certain length to achieve masking of the C-terminal domain.

### ΔRep2 and Scr sporozoites have decreased infectivity for hepatocytes

To assess the capacity of CSP repeat mutant salivary gland sporozoites to infect mice, we intravenously inoculated sporozoites into mice and quantified relative parasite abundance in the liver using RT-qPCR (Bruna-Romero et al., 2001). While ΔRep1 and ΔRep3 sporozoites had similar liver stage parasite burdens as controls, infection with ΔRep2 and Scr sporozoites resulted in significantly lower liver parasite burden (Figure 3A). To determine whether the defects in ΔRep2 and Scr infectivity were due to defects in invasion or development, we performed *in vitro* hepatocyte infection assays with Hepa1-6 cells and Scr sporozoites, as ΔRep2 salivary gland sporozoite yields were too low for these assays. Infection of hepatocytes with the same number of control or Scr sporozoites resulted in fewer Scr exoerythrocytic forms (EEFs) compared to controls (Figure 3B). However, Scr EEFs did not have defects in EEF development, as determined by EEF size (Figure 3B), suggesting that Scr sporozoites are defective in entering hepatocytes but once inside, they develop normally. Defective infection of hepatocytes by both ΔRep2 and Scr sporozoites was sufficient to delay the initiation of blood stage infection (pre-patency period) (Figure 3C).

**Figure 3:**
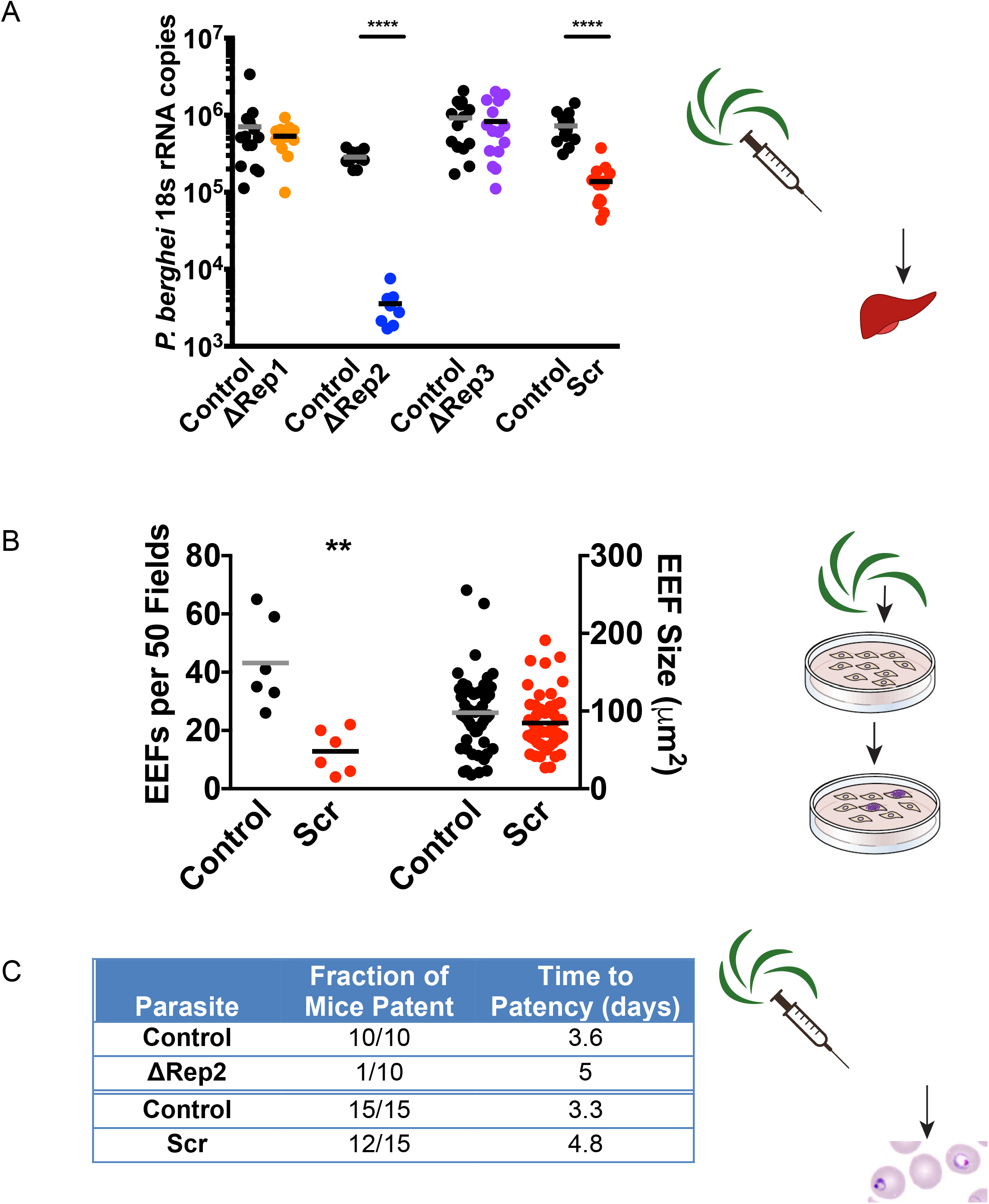
ΔRep2 and Scr sporozoites have decreased infectivity in the mammalian host. **(A)** Sporozoite infection assays in mice. 5000 control (WT-GFP) or CSP repeat mutant sporozoites were inoculated intravenously into C57Bl/6 mice and 40 hrs later, parasite liver burden was determined by RT-qPCR using primers specific *P. b*erghei 18s RNA. Shown are pooled data from 2 to 3 experiments with 5 mice per group in each experiment. Kruskal-Wallis p<0.0001 followed Mann-Whitney-Wilcoxon ****p<0.0001. **(B)** Development of exoerythrocytic stages (EEFs*) in vitro*. Control or Scr mutant sporozoites were added to Hepa1-6 cells and 24 hrs later EEFs were fixed and stained with a-HSP70. Shown are the numbers of EEFs per 50 fields and EEF size at 48 hours post-infection from two independent experiments with 3 replicates per experiment. Welch’s t-test **p<0.005. **(C)** Initiation of blood stage infection by ΔRep2 and Scr mutant sporozoites. 5,000 sporozoites were inoculated intravenously into mice and mice were monitored by blood smear from day 3 to day 9 post-infection. Shown are the number of mice that became patent and the average day of patency for mice with blood stage infections. Data are pooled from 2 to 3 experiments.

### ΔRep2 and Scr sporozoites have gliding motility deficits

ΔRep2 and Scr sporozoites exhibited a diminished capacity to infect two very different tissues: the salivary glands of mosquitoes and the liver of mice. Lack of tissue specificity in the infectivity phenotype of the CSP repeat mutants suggests these phenotypes are not the result of interrupted receptor-ligand interactions, but due to a feature required of sporozoites at both stages. Previous studies have demonstrated that sporozoites with motility defects cannot invade salivary glands or hepatocytes efficiently (Ejigiri et al., 2012; Sultan et al., 1997). Therefore, we analyzed the motility phenotypes of the CSP repeat mutants using several *in vitro* assays.

We began with live gliding assays, recording salivary gland sporozoite motility on glass coverslips for 2.5 minutes. Approximately 90% of sporozoites were attached and motile across the studied lines (Supplementary Figure 5). We categorized sporozoite motility according to previously observed motility phenotypes: continuous circular gliding, patch gliding, or attached waving (Vanderberg, 1974; Hegge et al., 2010). Examples of each motility type are shown in Figure 4A and in Videos S1-3. Circular gliding, in which sporozoites move in continuous circles, is the predominant form of motility observed in wild-type parasites and is an indicator of fully mature and infectious sporozoites (Vanderberg, 1974) (Video S1). Patch gliding sporozoites move back and forth over one area and do not demonstrate continuous forward motility. Patch gliding is rarely observed in wild-type salivary gland sporozoites, though it is frequently observed in midgut sporozoites and in mutants such as the TRAP KO parasite, where parasite adhesion is defective (Sultan et al., 1997; Kappe et al., 1999) (Video S2).These data have lead to the hypothesis that patch gliding is a step in the maturation of the motility machinery (Münter et al., 2009). Waving parasites are attached at one end and flex the other end (Vanderberg, 1974; Hegge et al., 2010) (Video S3). We found that like controls, the majority of ΔRep1 and ΔRep3 sporozoites engaged in circular gliding, with a small minority patch gliding or waving (Figure 4B). In contrast, ΔRep2 and Scr sporozoites had a significantly lower proportion of sporozoites engaged in circular gliding (12% ΔRep2 and 27% Scr), while the majority of these mutants were either patch gliding, (39% ΔRep2 and 46% Scr) or waving (41% ΔRep2 and 25% Scr).

**Figure 4:**
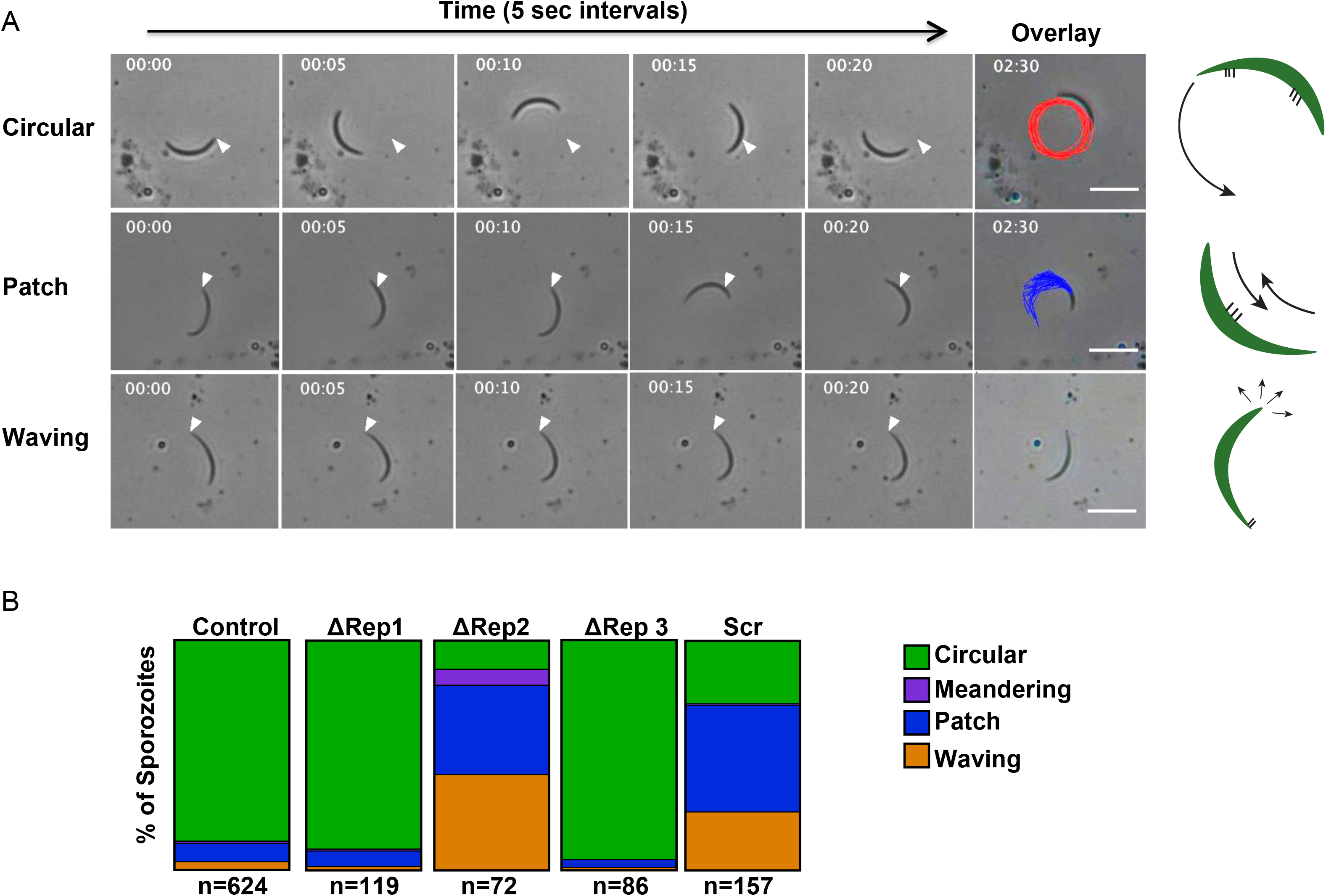
ΔRep2 and Scr sporozoites exhibit decreased productive gliding motility. CSP repeat mutants and control sporozoites were added to glass coverslips and their motility was recorded in 2 min movies and analyzed. **(A)** Examples of the three predominate classes of motility behavior observed: Shown are stills from movies in which sporozoites exhibit circular gliding (top), patch gliding (middle), and waving (bottom) with the tracking overlay from the entire movie in the final column. White arrow indicates the position of the sporozoite’s posterior end at the beginning of the movie. Scale bars = 10 µm. Cartoons on the right hand side show sporozoites in green, adhesion sites in thick black lines, and direction of motility (arrows). **(B)** Quantification of motility types for each mutant. Percent of bar occupied by a given motility behavior corresponds to percent of sporozoites exhibiting that behavior. Assays with mutant sporozoites were always done side-by-side with WT-GFP controls. At least three experiments per mutant were done. Total number of each genotype analyzed is indicated below each bar. There were statistically significant differences between the proportion of SCR and ΔRep2 sporozoites exhibiting circular gliding, patch gliding, and waving when compared to control sporozoites (Z-test for proportions, p<0.00001). There was also a significant difference in the proportion of meandering sporozoites between ΔRep2 and control sporozoites (Z-test for proportions, p<0.001).

(Figure 4B). These differences in motility phenotypes between the ΔRep2 and Scr sporozoites and controls were statistically significant (Z-test for proportions, p<0.00001).

### ΔRep2 and Scr sporozoites are slower circular gliders, but faster patch gliders

Substrate based gliding motility of sporozoites requires the cyclic formation and dissolution of adhesion sites (Münter et al., 2009; Hegge et al., 2010; Ejigiri et al., 2012). Adhesion cycles are comprised of the slow development of adhesion sites followed by fast movement over these adhesions (Münter et al., 2009) and can be observed in the instantaneous speed plots of gliding sporozoites (Supplementary Figure 6). To better understand the motility defects in ΔRep2 and Scr sporozoites, we analyzed these instantaneous speeds plots to determine how much time each mutant spent at a given speed. To do this we combined the data from all movies, extrapolated the speed between every frame of each movie and plotted the frequency of speed distributions. When data from circular gliding and patch gliding sporozoites were combined, ΔRep2 and Scr sporozoites were observed to move less often and at slower speeds (Figure 5A left panel).

**Figure 5:**
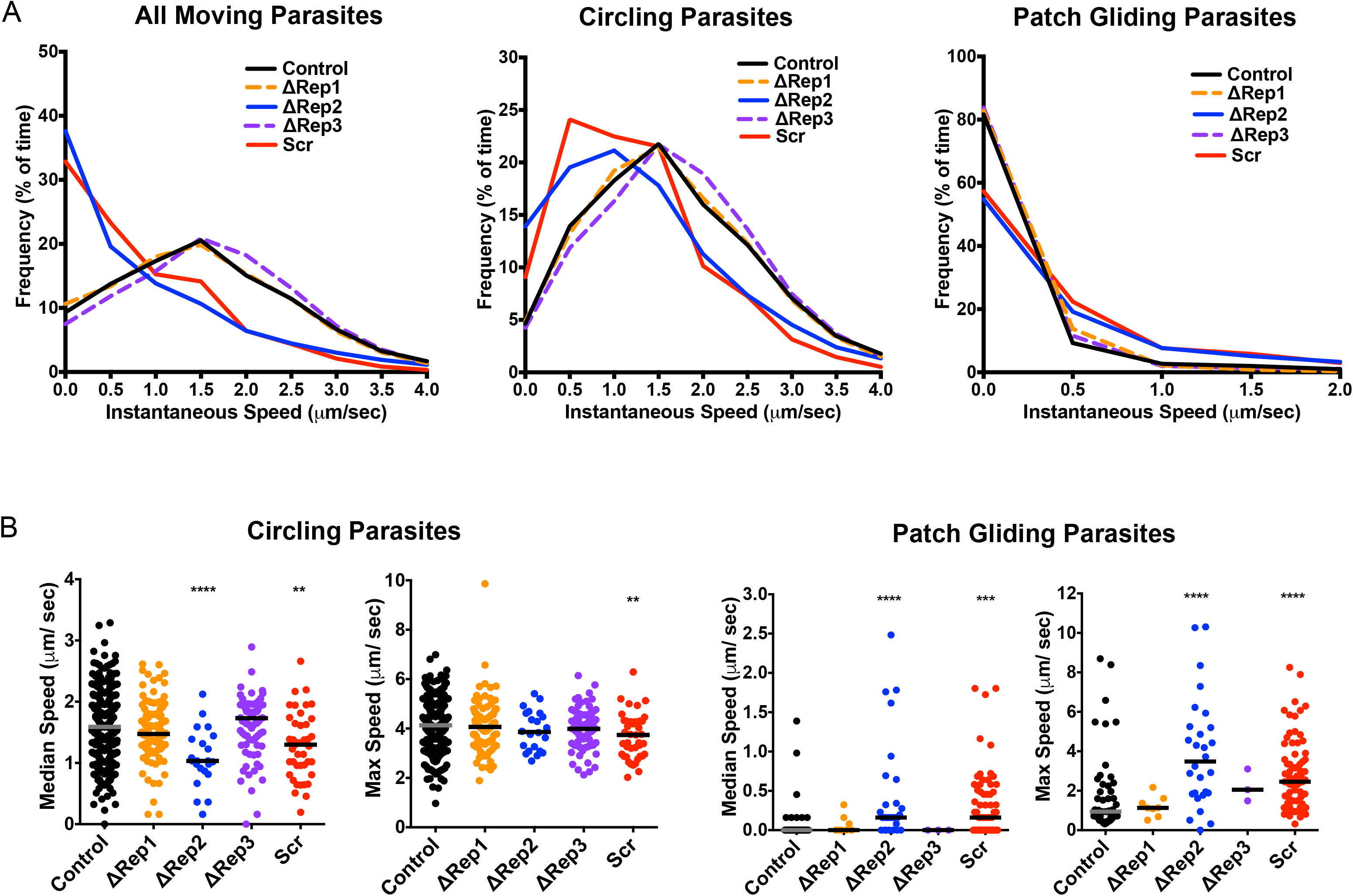
Instantaneous speed measurements demonstrate that ΔRep2 and Scr have decreased speed during circular gliding but faster patch gliding speeds. CSP repeat mutants and control sporozoites were added to glass coverslips and their motility was recorded in 2.5 min movies. **(A)** Frequency distributions of instantaneous speeds recorded for mutant and control sporozoites that were either circling or patch gliding (left), only circling (center) and only patch gliding (right). Instantaneous speed plots were pooled to determine the percent of total time that each population of sporozoites spent in each 0.5 µm per second speed bin. **(B)** Graphs show median and maximum speeds for each sporozoite recorded for circling sporozoites (left) and patch gliding sporozoites (right). Each mutant was analyzed from at least three independent mosquito cycles and at least 20 movies were collected per experiment. Controls are pooled from all experiments. For median speed of circling parasites: One-way ANOVA p<0.01 followed by Dunnett’s test ****p<0.0001, ** p<0.01. For all other data sets, Kruskal-Wallis p<0.0001 followed by Mann-Whitney- Wilcoxon **p<0.01,***p<0.0005, ****p<0.0001. Number of circular gliding sporozoites analyzed for each parasite line are as follows: Control=559, ΔRep1=108, ΔRep2=22, ΔRep3=82, Scr=43. Number of patch gliding sporozoites analyzed for each parasite line are as follows: Control=50, ΔRep1=8, ΔRep2=28, ΔRep3=3, Scr=73.

To look at this in more detail, we separately analyzed the instantaneous speed plots of circular and patch gliding parasites. We found that circular gliding ΔRep2 and Scr sporozoites spent more time not moving or moving at ≤ 0.5 µm/sec (Figure 5A middle panel) compared to controls or ΔRep1 and ΔRep3 sporozoites. Since circular gliding ΔRep2 and Scr sporozoites spent more time moving at slower speeds, it would be expected that their median speed, over the entire recorded movie, would be slower and indeed that is what we observed (Figure 5B left panel). Nonetheless, no severe defects in maximum speed were observed (Figure 5B left panel), suggesting that the motor is still functional in these mutants. In contrast to the circular gliding sporozoites, instantaneous speed distributions of patch gliding ΔRep2 and Scr parasites showed that these mutants exhibited faster speeds than control or ΔRep1 and ΔRep3 patch gliders and spent more time moving (Figure 5A right panel). These data are consistent with the increased median and maximum speed of ΔRep2 and Scr patch gliding parasites, both of which were higher than controls, ΔRep1 and ΔRep3 sporozoites (Figure 5B right panel), further suggesting that the internal motility machinery is intact.

### Adhesion site dynamics in ΔRep2 and Scr sporozoites

Gliding motility involves the formation and rapid turnover of adhesion sites (Münter et al., 2009; Hegge et al., 2010) on the sporozoite surface. To better understand the gliding defects in our CSP repeat mutants, we used reflection interference contrast microscopy (RICM) to observe adhesion site assembly and disassembly over time. RICM imaging phase-shifts the light reflected closest to the coverslip such that cell membranes closely opposed to the coverslip appear dark and can be distinguished from the rest of the cell (Weber, 2003). A previous study of circular gliding sporozoites found that an adhesion site first forms at the anterior end of the sporozoite, grows (assembles), sometimes becoming as large as the full length of the sporozoite, and finally disassembles, becoming smaller at the rear end of the sporozoite while the formation of another adhesion site at the anterior end of the sporozoite occurs, thus continuing the cycle (Münter et al., 2009) (Figure 6A and Video S4). In contrast, when patch gliding sporozoites are imaged by RICM, one observes a small site of adhesion over which the sporozoite repeatedly moves back and forth, unable to either grow this adhesion site or to form a second adhesion site (Figure 6A and Video S5). Thus, the high proportion of patch gliding ΔRep2 and Scr sporozoites suggests that these mutants have altered adhesion site dynamics.

**Figure 6:**
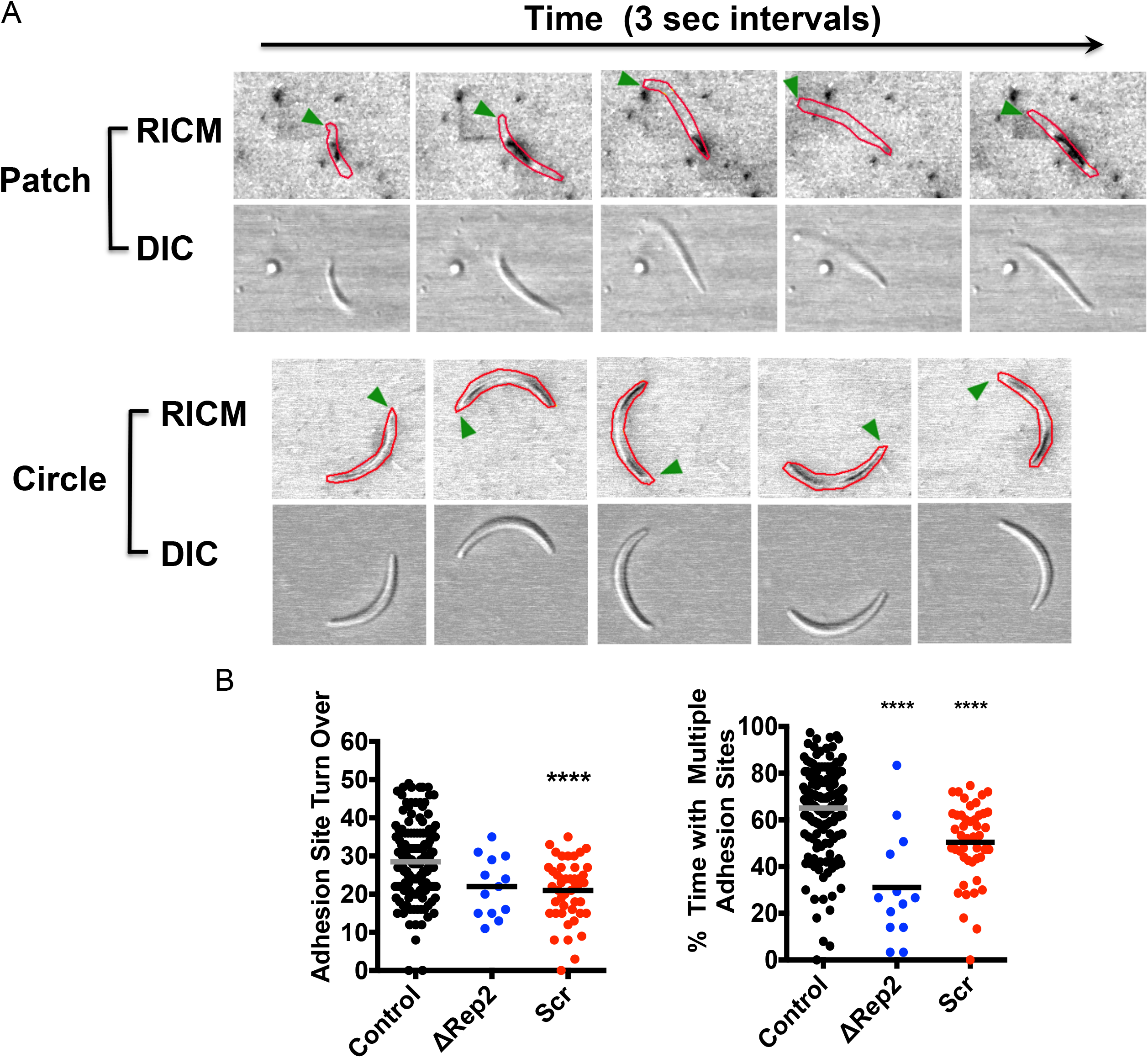
**ΔRep2 and Scr sporozoites have defects in adhesion site formation and turnover**. Sporozoites were added to glass coverslips and imaged by reflection interference contrast microscopy (RICM) to record adhesion site formation and turnover. **(A)** Sequential stills from RICM imaging of patch gliding and circular gliding sporozoites illustrate adhesion site dynamics during each motility type. RICM images below show adhesion sites with sporozoites outlined in red and green arrowheads indicating the same end of patch gliders and the anterior end of circular gliders. Corresponding DIC images of sporozoites are shown below. **(B)** Adhesion site formation and turnover of circular gliding control, ΔRep2, and Scr sporozoites. RICM movies of sporozoites were analyzed to quantify the frequency with which adhesion sites move from the sporozoite’s anterior to posterior end in 2.5 minutes of circular gliding and the frequency with which gliding sporozoites have multiple adhesion sites. One-way ANOVA p<0.0001 followed by Dunnett’s test ****p<0.0001. Data are pooled from 3 or more independent experiments with the following number of parasites for each line: Control=178, ΔRep2=13, Scr=46.

Since our instantaneous speed analysis of ΔRep2 and Scr circular gliders demonstrated that they spend more time not moving or moving slowly, we used RICM to visualize adhesion site turnover in the small percentage of ΔRep2 and Scr mutants that were engaged in circular gliding. We defined an adhesion turn over event as an adhesion site forming at the anterior end of the sporozoite, translocating to the posterior end, and finally disassembling (Figure 6A). When we quantified these events, we found fewer adhesion site turnover events per unit time in ΔRep2, and Scr mutants (Figure 6B). We then went on to dissect adhesion site assembly and disassembly. By observing RICM movies frame-by-frame we binned each time point as having multiple or single adhesion sites per sporozoite. Control sporozoites spend ∼70% of gliding time with multiple adhesion sites (Figure 6B). In contrast, circular gliding ΔRep2 and Scr sporozoites spend less time with multiple adhesion sites compared to controls (Figure 6B), suggesting an adhesion site dynamics defect.

### TRAP secretion does not explain adhesion site defects

TRAP (thombospondin-related anonymous protein) is a transmembrane protein that is critical for gliding motility (Sultan et al., 1997). It is secreted onto the sporozoite surface as sporozoites move, and functions to link the subpellicular actin-myosin motor to the extracellular substrate (Figure 7A) (Bosch et al., 2007; Ejigiri et al., 2012). Because of TRAP’s critical role in motility, we investigated whether it was secreted and distributed on the surface of ΔRep2 and Scr repeat mutants by performing immunofluorescence assays on gliding sporozoites using polyclonal antisera specific for TRAP. Control gliding sporozoites had relatively little TRAP staining on their surface, but substantial TRAP staining in the trails that gliding sporozoites leave behind. In contrast, we found that ΔRep2 and Scr sporozoites had a marked increase in detectable TRAP on the sporozoite surface, but TRAP was not detected in trails (Figure 7B). While 70-90% of Scr and ΔRep2 sporozoites do not engage in circular gliding (Figure 4), we would expect to see TRAP trails associated with the small percentage of mutant sporozoites that do engage in circular gliding. However, after counting 200 to 400 sporozoites we did not observe TRAP trails in these mutants. These data suggest that while TRAP is secreted onto the sporozoite surface in ΔRep2 and Scr sporozoites, it is not shed from the sporozoite surface, even by the small number of mutant sporozoites that do glide.

**Figure 7.**
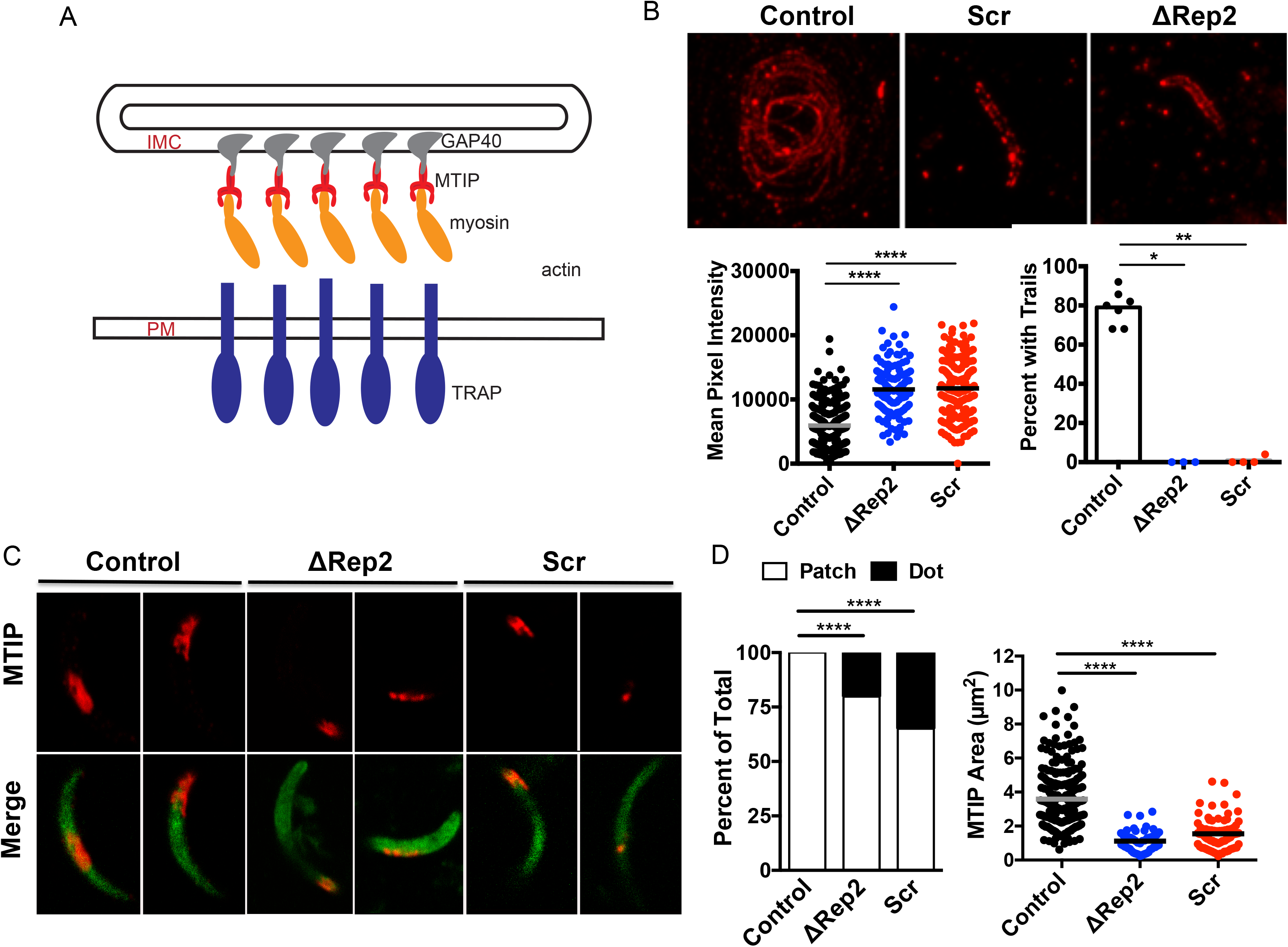
ΔRep2 and Scr sporozoites have defects in TRAP shedding and MTIP incorporation into adhesion sites. **(A)** Cartoon of the motor showing TRAP in the sporozoite plasma membrane (PM) with its cytoplasmic tail linked to the actin-myosin motor, which in turn is anchored to the inner membrane complex (IMC) via MTIP (myosin tail interacting protein) and various gliding-associated proteins (GAPs), of which GAP 40 is shown.**(B)** TRAP on circular gliding sporozoites was assessed by allowing them to glide on glass coverslips for an hour followed by staining for TRAP without permeabilization. Representative images of control, ΔRep2, and Scr sporozoites are shown in (A) and quantitative analysis of TRAP staining is shown in (B). Left panel: TRAP staining on the sporozoite surface was quantified by measuring mean pixel intensity of each sporozoite. One-way ANOVA p<0.0001 followed by Dunnett’s test ****p<0.0001. Right panel: Sporozoites with and without TRAP deposition in trails were counted and expressed as percent of total sporozoites. Kruskal-Wallis p<0.0005 followed by Mann-Whitney-Wilcoxon *p<0.05, **p<0.01. Data are pooled from 3 to 4 independent mosquito cycles with each mutant cycle having a paired control cycle. n = 20-100 sporozoites per experiment. Kruskal-Wallis p<0.0001. Mann-Whitney-Wilcoxon *p<0.05, **p<0.01 **(C&D)** MTIP staining on circular gliding sporozoites was assessed by allowing sporozoites to glide on glass coverslips, followed by fixation with paraformaldehyde and staining with antibodies specific for MTIP. Representative images of MTIP staining are shown in (C) and quantification of staining pattern and area occupied by MTIP are shown in (D). 20 to 50 sporozoites per line were identified by endogenous GFP and following this MTIP staining was marked and the area occupied by MTIP was quantified using Fiji. Staining pattern was manually counted with an example of the “dot” pattern shown in the second ΔRep2 and Scr images, and “patch” pattern examples shown in the control images and the first ΔRep2 and Scr images. Data are pooled from 2 to 3 experiments; Kruskal-Wallis p<0.0001; Mann-Whitney ****p<0.001.

The lack of TRAP in trails of the ΔRep2 and Scr repeat mutants, together with defects in adhesion site formation and turnover demonstrated by RICM analysis, prompted us to visualize adhesion sites in these mutants by staining for myosin tail domain interacting protein (MTIP) in gliding sporozoites. Previous studies have shown that gliding sporozoites have patches of MTIP on their surface that likely represent adhesion sites (Swearingen et al., 2016; Siden-Kiamos et al., 2020). As shown in Figure 7C&D, control sporozoites show patches of MTIP on their surface, which can be found on one end, in the middle, or at both ends of the sporozoite, similar to previously published results (Swearingen et al., 2016; Siden-Kiamos et al., 2020). In contrast, approximately 25% of ΔRep2 and 40% of Scr mutants had only one MTIP dot on their surface while the remainder had significantly smaller MTIP patches compared to controls (Figure 7C&D). The abnormal surface staining of both TRAP and MTIP on gliding ΔRep2 and Scr repeat mutant sporozoites, together with their inability to form and rapidly turnover adhesion sites, suggests the underlying problem in these mutants is on the sporozoite surface.

### The CSP Repeats have elastic properties

The finding that adhesion site formation on the sporozoite surface was impacted by mutations in the CSP repeats raised the possibility that the repeats have conserved biophysical properties that are important for motility. To date the CSP repeats have resisted structural analysis, likely due to a flexible and dynamic nature. Interestingly, CSP repeats have features in common with elastomeric proteins such as spider silk, titin, and abductin, which can crosslink to form a network (Tatham and Shewry, 2000). The elasticity of these proteins is defined by their ability to unfold and fold without energy loss or protein rupture (Tatham and Shewry, 2000). Additionally, elastic proteins are difficult to crystallize due to flexible repeats that are high in proline and glycine content and frequently have a ß- turn conformation. The CSP repeats have several of these properties: 1) they are high in proline or glycine content (Supplementary Table); 2) peptide structure studies have suggested a ß-turn conformation (Plassmeyer et al., 2009; Ghasparian et al., 2006; Fasman et al., 1990; Dyson et al., 1990; Verdini et al., 1991) and 3) they have proven difficult to crystalize.

To better understand the biophysical properties of the CSP repeats, we utilized optical tweezers combined with confocal single molecule fluorescence microscopy (Hohng et al., 2007) and performed single-molecule fluorescence-force spectroscopy on repeat and scrambled repeat peptides (Figure 8A) as previously performed for spider silk (Brenner et al., 2016; Grashoff et al., 2010). In these experiments, single-molecule FRET (Ha et al., 1996) signal is measured between two fluorophores, Cy3 (FRET donor) and Cy5 (FRET acceptor), flanking the peptide of interest while the peptide is under tension. One end of the peptide-fluorophore construct immobilized on a passivated glass coverslip while the other end is conjugated to a bead in an optical trap through DNA linkers (Figure 8A). Upon moving the microscope stage, force is applied to the peptide and peptide extension can be measured by changes in the FRET efficiency. As shown in Figure 8A, when the peptide is relaxed the FRET signal is high, as the Cy3 and Cy5 fluorophores are in proximity, while upon application of force, the fluorophores move apart as detected through reduction in FRET. FRET efficiency is restored to the high value when the peptide is relaxed through to the original low force. With elastic peptides that can stretch without heat dissipation, FRET vs force curves of the stretching and relaxation phases should overlap, which is indeed what we observed for a 25 amino acid CSP repeat peptide, (NANDPAPP)_3_ (Figure 8B). In contrast, the scrambled peptide had a relaxation phase with a lower FRET efficiency than during the stretching phase, indicating that the peptide does not rapidly recover its original compact state during force relaxation (Figure 8B). Such hysteresis indicates that the scrambled peptide behaves as an inelastic spring and that the elastic behavior observed for the CSP repeat peptide is a biophysical feature of its specific arrangement of amino acids. A similar elastic behavior was also observed for a longer 40 amino acid CSP repeat peptide (NANDPAPP)_5_ with the only difference being that the FRET efficiencies are lower due to the increased peptide length (Figure 8C). Finally, we converted FRET efficiency into the end-to-end distance of the peptide after correcting for the linkers between the fluorophores and the peptide, and found that the CSP repeats are linear springs, i.e. their extension increases linearly with force (Figure 8D). Compliance, defined as the extension change per unit force change, is larger for the longer peptide, further confirming that our experimental scheme indeed measures the mechanical properties of the CSP repeat peptides. The compliance values are about four-fold smaller than those measured from spider silk peptides, suggesting that the CSP repeat peptides are much stiffer. Together these data suggest that the CSP repeat region has properties of a linear-elastic spring and that the elasticity of the region is dependent on the repeated, primary amino acid sequence and the length of the repeats.

**Figure 8:**
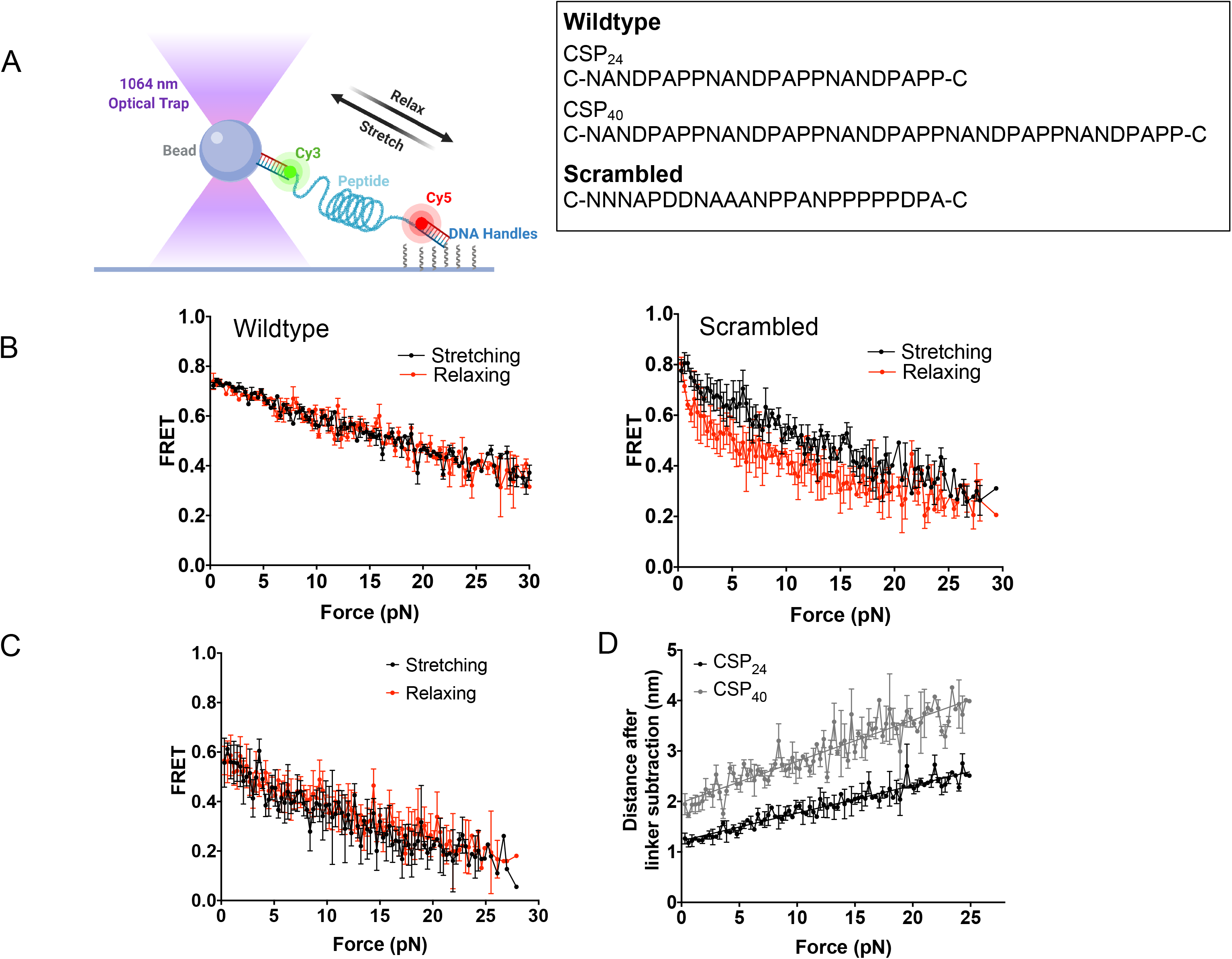
CSP Repeats have elastic properties. **(A)** Peptide sequences used in study and schematic of single-molecular force spectroscopy (SMFS). Wildtype and scrambled CSP repeat peptides were synthesized and prepared for single-molecule force spectroscopy studies with the addition of DNA handles and fluorophores as shown. SMFS set-up showing one end of the peptide conjugated to a DNA handle that then attaches to a bead caught in an optical trap and the other end of the peptide is immobilized on a glass slide. As the bead is pulled, it stretches the peptide, increasing the distance between the fluorophores, thus decreasing the FRET signal. Upon relaxation, the fluorophores are brought closer together and the FRET signal increases. **(B)** FRET signal during stretch (black) and relax (red) phases of wildtype peptide pulling n=22 and scrambled peptide pulling. n=16. Shown in are means +/- standard error. **(C)** A longer WT CSP repeat sequenced, CSP_40_, was prepared and tested in single-molecule fluorescence force spectroscopy studies. FRET signal during stretch (black) and relax (red) phases of wildtype peptide pulling. Error bars are SEM, n=22. **(D)** Wild type CSP repeats act as a linear spring independent of SMCC linker. Compliance (the distance the peptide stretched upon increasing force after linker subtraction) of the peptides is dependent on length and fits a linear fit model, a feature of linear springs. Two WT peptide lengths were tested: CSP_40_ and CSP_24_. Lines = liner fit model. R^2^ values for CSP_24_ = 0.93 and for CSP_40_=0.86. Error bars are SEM. n = 22 for both peptides.

Though a previous study using atomic force spectroscopy (AFM) approach also suggested an elastic CSP repeat region (Patra et al., 2017), the constructs used in that study had I27 domains from titin flanking the CSP repeats to provide a molecular fingerprint in the stress-strain curves so that the peaks associated with the CSP repeats could be identified. However, they found that the CSP repeats unfolded before the I27 bands, meaning that contributions of the I27 domains to the CSP elasticity data cannot be confidently excluded (Patra et al., 2017). As our constructs have no such domains and the compliance of the peptides are dependent on their length, we can confidently extrapolate the mechanical properties of the CSP repeats. Interestingly, the CSP repeats are substantially stiffer (∼4x) than the previously characterized spider silk proteins, making them an interesting potential molecular tension sensing tool (Brenner et al., 2016).

## Discussion

In this study we investigated the structural and functional properties of the CSP repeats by generating mutant parasites with altered repeats and using single molecule mechanical measurements. These orthogonal approaches both demonstrate that the length and organization of the repeats into repetitive sequence blocks is essential to their functional properties, putting to rest the long-held belief that the CSP repeats are a non-functional linker, connecting two flanking domains, and without conserved structural properties (Schofield, 1990).

The repeat mutants generated in the current study have normal sporozoite development in the mosquito, indicating that the mutations we introduced do not lead to gross misfolding of CSP and allowing for downstream functional analysis of the repeats in salivary gland sporozoites. Though a role for the repeats in gliding motility has been hypothesized based on the finding that high titers of anti-repeat antibodies and Fab monomers can immobilize sporozoites (Stewart et al., 1986), the mechanism for such a role has not been elucidated. Indeed, since CSP is not a transmembrane protein and therefore cannot link to the motor, it has not been considered in the majority of studies probing the mechanism(s) by which sporozoites move. Here we provide several lines of evidence demonstrating that the CSP repeats have a critical role in sporozoite motility. The two mutants with significant deficits in motility, ΔRep2 and Scr, demonstrate that both length and organization into repetitive sequence blocks are critical to CSP repeat function. These mutants have a diminished capacity for productive motility, with an enhanced proportion of sporozoites patch gliding or waving, suggesting a problem with secondary and/or large adhesion site formation. This was confirmed with RICM imaging of gliding sporozoites, which demonstrated severe adhesion site formation defects in patch-gliders, a gliding phenotype rarely observed in wild-type sporozoites and commonly observed in ΔRep2 and Scr sporozoites. Furthermore, the small percentage of circular gliding ΔRep2 and Scr sporozoites demonstrated defects in adhesion site formation and turnover.

Three observations suggest that the defect in adhesion site formation in in ΔRep2 and Scr sporozoites is due to dysfunction on the sporozoite surface and not to problems with the internal motility: 1) patch gliding ΔRep2 and Scr salivary gland sporozoites move faster than patch gliding control sporozoites; 2) circular gliding ΔRep2 and Scr sporozoites turn over their adhesion sites at a slower rate, yet reach maximum speeds comparable that of controls; 3) TRAP is secreted onto the parasite surface in ΔRep2 and Scr sporozoites, suggesting the gliding defects in these mutants is downstream of TRAP secretion. Since motility in both patch gliding and circular gliding are dependent upon actin polymerization (Münter et al., 2009), these data suggest that ΔRep2 and Scr repeat mutants have fully developed their internal motility machinery.

TRAP, the dominant adhesin during sporozoite motility, must be shed from the surface to allow the parasite to disengage from adhesive interactions and move forward, a process that requires the activity of plasma membrane rhomboid proteases (Ejigiri et al., 2012; Shen et al., 2014; Buguliskis et al., 2010). Our finding of increased TRAP on the surface of ΔRep2 and Scr sporozoites is similar to what was observed in sporozoites in which the rhomboid cleavage site of TRAP was mutated (Ejigiri et al., 2012). However, in contrast to the CSP repeat mutants, the TRAP cleavage site mutants form adhesion sites from which they have trouble disengaging, leaving thick TRAP trails in their wake (Ejigiri et al., 2012). While the majority of rhomboid cleavage site mutant sporozoites engage in productive motility, only few of the ΔRep2 and Scr mutants are capable of productive motility, suggesting that what happens to TRAP after its secretion onto the sporozoite surface differs between these two sets of mutants. In the rhomboid cleavage site mutants, adhesion sites are formed but cannot be removed. In contrast, in the CSP repeat mutants, the formation of mature adhesion sites rarely occurs, as evidenced by the increased proportion of patch gliders and decreased adhesion site turnover in the small proportion of circular gliders in ΔRep2 and Scr mutants. The lack of TRAP trails in the CSP repeat mutants suggest that without a proper adhesion site, the rhomboid protease may not have access to TRAP. Though there may be some spots of TRAP in the trails that we cannot discern because of the relatively high background of our polyclonal antisera, it is clear that TRAP is not shed from the surface of the ΔRep2 and Scr mutants in the same manner as controls.

Visualization of MTIP patches on the surface of ΔRep2 and Scr mutants further supports the finding that adhesion site formation is defective in these mutants. Unfortunately, it has been difficult to visualize TRAP during gliding motility: GFP-tagging of TRAP results in non-motile sporozoites (Kehrer et al., 2016) and fixation of gliding sporozoites does not reproducibly demonstrate TRAP in patches, likely due to its movement during the fixation procedure. However, we and others have demonstrated that components of the motor that interact with the inner membrane complex, such as MTIP and GAP40, are stable during fixation and can be visualized in patches on the gliding sporozoite’s surface (Swearingen et al., 2016; Siden-Kiamos et al., 2020). MTIP staining on gliding control and mutant sporozoites showed that in contrast to controls, MTIP was less frequently observed in patches and when present, the patches were significantly smaller on ΔRep2 and Scr sporozoites, supporting the hypothesis that proper adhesion sites are not formed on the surface of these mutants. Taken together, our data suggest that the motility defects observed in the ΔRep2 and Scr repeat mutants are due to abnormal adhesion site formation and dynamics, resulting from alterations in the organization of CSP on the sporozoite surface. We hypothesize that the CSP repeats enable the normal formation and turnover of adhesion sites by providing a cohesive environment in which adhesion sites can form and be maintained (Figure 9). This is supported by a previous study in which optical tweezers were used to probe the sporozoite surface and found spatially segregated functional effects on motility suggesting that there are distinct cohesive capacities on the sporozoite surface (Hegge et al., 2012).

**Figure 9:**
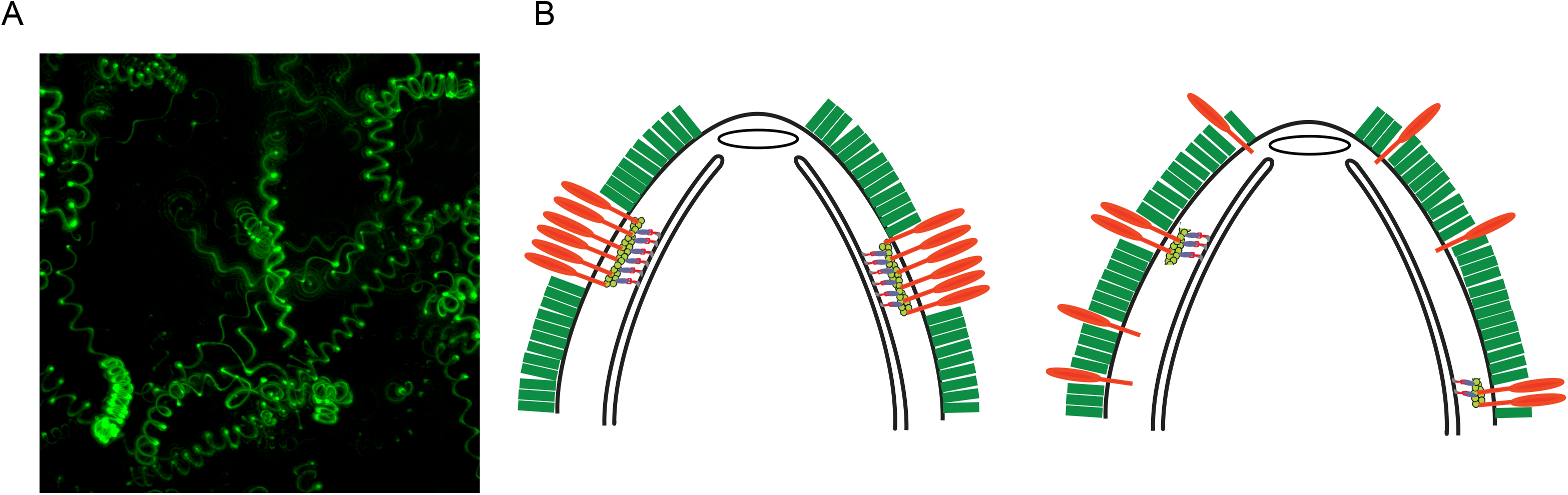
Sporozoite deformability in 3-D matrix and model of adhesion site formation in wildtype and mutant sporozoites. **(A)** Maximum projection of a one-minute movie showing tracks of wild-type *P. berghei* sporozoites as they move in 3-dimensions through matrigel. Note the different slopes of the helices. **(B)** Cartoon of the anterior portion of a wildtype (left) and CSP repeat mutant (right) sporozoite, with CSP (green) on the plasma membrane and TRAP (orange), forming a proper adhesion site in wildtype sporozoites (left), or being distributed throughout the mass of surface CSP unable to form an adhesion site (right). Underneath the plasma membrane a simplified motor complex embedded in the inner membrane complex is shown.

Interestingly, deletion of 25% of the repeat region, whether it be in the N-terminal (ΔRep1) or the C-terminal (ΔRep3) portion of the repeats, does not significantly impact function. In contrast, deletion of 50% of the repeat region (ΔRep2) leads to the motility defects outlined above and increased TSR- domain exposure. As shown in Figure 1A, the central repeat region links the N- and C-terminal domains, with previous studies demonstrating that the TSR domain is masked in salivary gland sporozoites until reaching the liver (Coppi et al., 2011; Hopp et al., 2015). Thus, our data suggests that there is a length requirement for masking of the TSR. In field isolates, the number of CSP repeats can vary up to ∼12% of the repeat length and modeling of repeat region length has suggested that length may be important for stabilizing repeat region structure (Gandhi et al., 2014; Escalante et al., 2002). The elastic properties of the CSP repeats could play a role in the masking of the TSR domain with the 50% truncation in ΔRep2 not allowing for sufficient stretch to mask this domain.

Our biophysical studies demonstrating that the CSP repeats have elastic properties and behave as a stiff, linear spring are concordant with the motility defects of the repeat mutants. Both normal gliding motility and the biophysical properties of the repeats are dependent upon length and their organization into repetitive blocks. While we do not directly demonstrate a connection between the elastomeric properties of the repeats and their role in motility, we hypothesize that these properties are important in the phenotypes we observed in our mutants. Previous studies have found that proteins with elastomeric properties interact with one another to form higher order structures that confer new mechanical and structural properties critically important to their function (Tatham and Shewry, 2000). Thus, these properties may be important for both the organization of CSP on the sporozoite’s surface, enabling the cohesion of the repeats and allowing for the formation and organization of proper adhesion sites, and for the regulated exposure of the TSR domain. Studies exploring whether the elastomeric properties of the repeats orchestrate the separation of CSP from adhesion sites on the sporozoite surface would be of particular interest. It is also possible that the elastic properties of the repeats modulate the dramatic changes in sporozoite curvature as they migrate through three-dimensional matrices *in vivo*. Alternatively, they might enable the parasites to keep substrate contact as the sporozoites bend upon interaction with the environment. 3D-projections of wildtype sporozoites migrating through matrigel (Figure 9) illustrate the curvature changes they experience during motility *in vivo*. Identifying the strength of the molecular forces experienced by the sporozoite, and its surface proteins, during adhesion and migration could elucidate the functional importance of an elastic repeat region.

In conclusion, the finding that mutations to the CSP repeats cause organizational changes on the sporozoite’s surface sufficient to impact adhesion site formation supports a long-held, but unproven, idea that CSP forms a higher order structure on the sporozoite surface (Ozaki et al., 1983). While this notion is supported by the dense packing of CSP on the sporozoite surface and the observation that CSP is shed as an intact coat upon cross-linking with antibody (Vanderberg et al., 1969), little progress has been made on defining the structure of CSP. Our current model is that CSP, by forming a cohesive entity, allows the components of adhesion sites to assemble in proximity to one another rather than be randomly distributed in a sea of CSP. Overall, our findings demonstrate that the CSP repeats have an important function in gliding motility and studies exploiting this function should aid the malaria vaccine effort focused on the sporozoite stage of the parasite.

## Methods

### Ethics Statement

All animal work was conducted in accordance with the recommendations in the Guide for the Care and Use of Laboratory Animals from the National Institutes of Health. This work was approved by the Johns Hopkins University Animal Care and Use Committee (protocols RA110608, M011H467 and M014H363), which is fully accredited by the Association for the Assessment and Accreditation of Laboratory Animal Care.

### Antibodies

Anti-C terminal sera were previously generated against a 100 amino acid peptide representing the C-terminal third of CSP (Coppi et al., 2005). mAb 3D11 is specific for the *P. berghei* CSP repeats (Yoshida et al., 1980), and mAb 2E6 is specific for *Plasmodium* Hsp70 (Tsuji et al., 1994). Antisera against the *P. berghei* TRAP repeats is specific for the C-terminal repeat region of TRAP (Ejigiri et al., 2012). Antisera to a predicted disordered region immediately C-terminal of the CSP repeat region and upstream of the TSR (anti-C-dis), was generated in a New Zealand White rabbit using the peptide, CDDSYIPSAEKILEFVKQIRDSITE, synthesized and purified by RS Synthesis (Louisville, KY). The peptide was conjugated to keyhole limpet hemacyanin and the rabbit was immunized once and boosted 3 times at ≥1 month intervals as previously outlined (Harlow and Lane, 1988).

### Plasmid Generation

Mutant *csp* repeat region oligonucleotides were commercially synthesized by Genscript (Piscataway, NJ) and provided in the pUC57 plasmid. Repeat sequences were digested from the pUC57 plasmid using HindIII-HF and SexAI and ligated into an intermediate CSP vector in Bluescript SK, containing the KpnI to PacI sequence from the *csp* locus which includes 500 bp of 5’utr and the full length *csp* sequence. The *csp* sequence in this vector was engineered to have HindIII-HF and SexAI restriction sites flanking the repeats. Once the mutant repeat sequences were ligated into the intermediate CSP vector, a KpnI and PacI digest was performed and this fragment was ligated into the previously described transfection vector, pCSRep, altered to have an additional 1 kb of 3’ UTR in order to minimize correction of the repeat mutations (Coppi et al., 2011; Ferguson et al., 2014) (Supplementary Figure 1A).

### Generation and Verification of *P. berghei* CSP Repeat Mutants

Recombinant parasites were generated in the *P. berghei* ANKA 507 cl1 line which expresses GFP under the *eef1α* promoter at the 230p locus (WT-GFP) (Janse et al., 2006). Recombinant parasites were generated by double homologous recombination in which the native *csp* locus was replaced by a mutant copy of *csp* with its control elements and an upstream selection cassette (Supplementary Figure 1). pCSRep was digested with XhoI and KasI and 5-10 µg of DNA was electroporated into schizonts which were injected intravenously into a Swiss Webster mouse, selected with pyrimethamine, and cloned by limiting dilution in Swiss Webster mice as previously described (Janse et al., 2006). Repeat mutant genotypes were confirmed by PCR and sequencing. 5’ integration was confirmed using primers DP1 (5’-AATGAGACTATCCCTAAGGG-3’) and DP2 (5’- TAATTATATGTTATTTTATTTCCAC-3’), 3’ integration was confirmed using primers P6 (5’- TGATTCATAAAT-AGTTGGACTTGATTT-3’) and P9 (5’- TCGAAATGGGCGCTGAC AAGAA-3’) and amplification of the *csp* locus was used to confirm repeat mutations using primers P3 (5’- CCATTTTAGTTGTAGCGTCACTTTT-3’) and P4 (5’-ACAAATCCTAATGAATTGCTTACA-3’).

Absence of WT parasites in mutant clones was confirmed using size shift PCR. P14 (5’- CGTGCATTTTGTGTCCTCATGTTGC-3’), binds in the 5’ UTR of *csp*, and P17 (5’-GCTCGTTTAAGTTCCTTTGGGCTTGG-3’), binds the 5’ end of *csp*. This primer pair resulted in a 1.46 kb product in WT parasites and a 4.8 kb product in recombinant parasites (Supplementary Figure 1E). After PCR confirmation, *csp* was amplified by PCR and sent for sequencing.

### Mosquito Infection and Sporozoite Isolation

*Anopheles stephensi* mosquitoes were reared in the insectary at the Johns Hopkins Bloomberg School of Public Health and infected with control and mutant parasite lines by feeding on anesthetized Swiss Webster mice as previously outlined (Vanderberg and Gwadz, 1980). Salivary gland sporozoites were harvested from salivary glands dissected in Lebowitz media (L-15, Gibco #11415-064) 21-25 days after mosquito feeds. Dissected salivary glands were briefly spun to remove the media and add fresh L-15 prior to releasing sporozoites by homogenization. Sporozoite preps were then centrifuged at 4°C for 4 minutes at 100xg and sporozoites were collected from the supernatant.

### Quantification of Midgut and Salivary Gland Infections

Oocyst numbers were assessed at 10-14 days post blood meal. Mosquito midguts were dissected, mounted on a glass slide and imaged with a 4x objective on a Nikon Eclipse E600 upright microscope and counted using the GFP fluorescence of parasites. Quantification of sporozoites in the midgut, hemolymph, and salivary glands was performed on mosquitoes asorted by GFP fluorescence. Twenty midguts and salivary glands were dissected on day 16 or 21 post bloodmeal, respectively, homogenized to release sporozoites which were then counted on a hemocytometer. Hemolymph was collected by perfusion of the mosquito abdomen on day 18 post blood meal and sporozoites were counted as above. For all sporozoite counts, the total number of collected sporozoites was divided by the number of mosquitoes to calculate the average number of sporozoites/mosquito. Salivary glands of ΔRep2 and Scr infected mosquitoes were imaged to confirm that sporozoites invaded the glands. Salivary glands were cleanly dissected from infected mosquitoes placed into 5 µl of L-15 on a MaTek well (MaTek #P35G-0-14C) followed by the addition of 5 µl of matrigel (BD #356231). The matrigel was allowed to polymerize at room temperature for ∼10 minutes and Z-stacks of salivary glands were acquired using a 3i spinning disk confocal microscope (Zeiss Axio Observer Z1 microscope with Yokogawa CSU222 spinning disk), with a 472 nm laser to observe GFP expressing sporozoites and DIC to observe the gland architecture.

### Western Blots

Anti-Cdis, which targets a predicted disordered region between the repeats and TSR of CSP was used for CSP quantification by western blot since antibodies specific for the repeat region could not be used in our mutants. For ΔRep1, ΔRep3, and RCon lines, salivary gland sporozoites were isolated as described and for ΔRep2 and Scr lines, hemolymph sporozoites were used, as low parasite abundance in the salivary glands prevented clean western blots. Sporozoites were pelleted, resuspended in sample buffer (0.125 M Tris-HCl, 20% glycerol, 4% SDS, 0.002% bromophenol blue, 50 mM DTT, and 1x protease inhibitors (Roche #11-836-153-001) at a concentration of 10^4^ sporozoites/µl, and 100,000 sporozoites were loaded per well. Samples were run on a 10% SDS-PAGE gel and transferred to a nitrocellulose membrane. CSP was detected using anti-Cdis polyclonal sera (1:100) and TRAP, used as a loading control, was detected with anti-TRAP polyclonal sera (1:500) (Ejigiri et al., 2012). Western blots were developed with goat anti-rabbit HRP antibody (1:10,000; GE Healthcare #NA934V) and ECL reagent (GE Healthcare #RPN2106).

### Immunofluorescence Assays (IFAs)

Salivary gland sporozoites were isolated from mosquitoes as described above and spun onto 12 mm coverslips in a 24-well plate at 300xg for 3 minutes at 4° C with low acceleration and no brake. Sporozoites were fixed for 1 hour with 4% PFA at room temperature, washed, blocked with 1% BSA/PBS, and incubated at 37°C with anti C-terminal sera (1:100) in 1% BSA/PBS. When indicated, 0.1% saponin was incorporated into primary antibody dilutions to break intra-molecular interactions on the surface of the sporozoite. For TRAP staining, a “gliding IFA” was performed. After sporozoites were spun onto coverslips, media was exchanged for 2% BSA/L-15, pH 7.4 and incubated for 1 hour at 37°C with 5% CO_2_. After this they were fixed as above, blocked in 1% BSA/5% goat serum/PBS and incubated with anti-TRAP repeat antisera (1:100) in 1% BSA/5% goat serum/PBS. After incubation with the appropriate secondary antibody coverslips were mounted with Prolong Gold and image acquisition was performed using a 100x objective on a Nikon Eclipse E600 upright microscope with a DS-Ri1 digital camera under identical acquisition settings for each treatment condition. Sporozoites were identified by phase-contrast, and then fluorescence imaging was performed to avoid bias acquisition. Intensity measurements were quantified using NIS Elements Br 3.2 software by using the “Auto ROI” function to identify the sporozoites in phase before switching to the fluorescence channel for intensity quantification. For MTIP and TRAP staining, freshly dissected sporozoites were spun onto 12 mm coverslips in a 24-well plate at 300g for 5 minutes and allowed to glide at 37°C for 30 minutes in 1% BSA in L15, pH7.4. Sporozoites were then fixed for 1 hour with 4% PFA at room temperature, blocked with 1% BSA/PBS for 30 minutes at room temperature and incubated with rabbit polyclonal anti-MTIP (1:500; Bergman et al., 2003) or anti-TRAP (1:100; [20]) in 1% BSA/PBS followed by detection with Alexa fluor 546 goat anti-rabbit secondary antibody (1:500, Invitrogen #A11010) in PBS. Samples were mounted in gold antifade mountant (Invitrogen, P36935) and image acquisition was performed using 100x objective with a 488 and 561 laser on a confocal microscopy (ZEISS, LSM800).

### Metabolic Labeling

Sporozoites were metabolically labeled and CSP was immunoprecipitated and detected as previously outlined (Coppi et al., 2005). Briefly, mosquito dissections were performed using DMEM without Cys/Met (Corning #17-204-C1) and sporozoites were metabolically labeled in DMEM without Cys/Met, 1% BSA and 400 µCi/ml l-[35S]Cys/Met (MP Biomedicals #0151006) for 45 minutes (ΔRep1) or 1.5 hr (ΔRep2) at 28°C. Labeled sporozoites were washed 3 times, lysed and CSP was immunoprecipitated with mAb 3D11 conjugated to agarose beads, eluted from beads with 0.1 M glycine pH 1.5 and 1% SDS and analyzed by SDS-PAGE followed by autoradiography.

### *In vivo* Sporozoite Infectivity Assays

5,000 sporozoites were injected intravenously into mice in 200 µL of cell culture media. For pre- patency experiments, blood smears were made from days 3-9 after inoculation of sporozoites, stained with Giemsa (Sigma-Aldrich #GS500), and screened for 5 minutes for the presence of parasites. For liver load experiments, mice were sacrificed 39 hours post sporozoite inoculation, livers were harvested, and RNA was extracted as previously outlined (Bruna-Romero et al., 2001). Parasite liver load was quantified by RT-qPCR using primers specific for *P. berghei* 18s rRNA and compared to a plasmid standard curve [19]. 4-5-week-old Swiss Webster mice (Taconic) were used for pre- patency experiments and 4-5-week-old C57BL/6 mice (Taconic) were used for liver load experiments.

### *In vitro* Hepatocyte Infection Assays

Hepa1-6 cells (ATCC #CRL-1830), a mouse hepatocyte cell line, were seeded on collagen I coated LabTek wells (Lab-Tek #17745) with 100,000 cells/well in 400 µl of cell culture media (DMEM, 10% fetal calf serum, and 2 mM gentamycin) 1 day before infection with sporozoites. 20,000 sporozoites were added to each well and centrifuged onto cells at 300xg for 3 minutes at room temperature with low acceleration and no brake. Sporozoites were then allowed to invade hepatocytes for 1 hour at 37°C with 5% CO2, after which the media was replaced and cultures were maintained with 1x penicillin-streptomycin (Gibco #15140-122). Media was changed twice daily and at 24- or 48-hours post-infection, hepatocytes were fixed with 4% PFA for 1 hour at room temperature. Cells were permeabilized with 100% methanol for 30 minutes at -20°C, followed by blocking with 1% BSA/PBS at 37°C for 1 hour. Parasites were stained with 10 µg/mL mAb 2E6 in 1% BSA/PBS for 1 hour at 37°C, followed by 30 minutes with 0.1% Tween/PBS at 37°C and 1 hour at 37°C with secondary antibody Alexa-488 goat anti-mouse (1:500) (Life Technologies #A11029) in 1% BSA/PBS before mounting with Prolong Gold with DAPI (Invitrogen #P36935). The number of EEFs per 50 fields at 24 hours post-infection was quantified for each well. For quantification of EEF size, the 2D area was measured using NIS Elements “Auto ROI” function on a Nikon Eclipse E600 microscope with a DS- Ri1 digital camera.

### *In vitro* Live Gliding Assays

Salivary gland sporozoites were dissected and a concentrated sporozoite suspension was mixed with an equal volume of 2% BSA (Sigma #A788) in L-15, pH 7.4 and incubated at 37°C for 5 minutes to activate sporozoites. Activated sporozoite suspension was placed between a glass slide or 22 mm X 50 mm cover glass and a 22 mm x 50 mm cover glass. Sporozoites were allowed to settle for 2 minutes before imaging. These assays were performed using a Nikon Eclipse E600 upright microscope using a 40x phase objective and imaged using phase contrast microscopy. Sporozoites were manually tracked using Fiji (http://fiji.sc). Sporozoite motility behavior was categorized as patch gliding if parasites moved over a single point without net displacement, as waving if sporozoites were attached without movement over the attachment site, and as circular gliders if they moved in circles, with displacement from the point of origin. Almost all sporozoites exhibited one form of motility during image acquisition. On the rare instances where sporozoites switched modalities, they were classified as the modality that is the most migratory (i.e. if a sporozoite did waving and circular gliding, it was classified as circular gliding). Sporozoite speed between each frame, instantaneous speed, was outputted using the Fiji “Manual Tracking” plugin. Frequency distributions were made by combining the instantaneous speed data points from all sporozoites within a group of interest (i.e. circular gliders). A frequency distribution was generated using 0.5 µm/sec speed bins and then plotted as a line graph for visualization.

### Reflection interference contrast microscopy (RICM)

RICM was performed using a LSM 780 inverted Zeiss AxioObserver with 780-Quasar confocal module and high-sensitivity gallium arsenide phosphide detectors (GaAsP) with a 63x (NA 1.4) oil objective and excitation with a 488-argon laser. A dichroic beam splitter (transmitted 80%, reflected 20%) was used to collect light emitted between 414-690 nm. All movies were acquired at 1 frame per second for 2.5 minutes. All movies were acquired at 1 frame per second for 2.5 minutes. For RICM analysis of adhesion sites, each frame was manually categorized as a sporozoite with a single or multiple adhesion sites. Percent of time with multiple adhesion sites was calculated by dividing the number of frames with multiple adhesion sites by the total number of frames (150) and multiplying by 100. Adhesion site turnover was determined by quantifying the number of adhesion sites per movie (2.5 minutes) that appeared at the anterior end and translocated to the posterior end of the sporozoite followed by complete disassembly.

### Single Molecule Fluorescence-Force Spectroscopy

To generate the peptide-oligomer constructs with Cy3 and Cy5 at either end, the protocol was adapted from a previous study (Brenner et al., 2016). Briefly, the DNA-oligonucleotide, ACCGCTGCCGTCGCTCCG with a 5’ amine modification, was incubated with 200x molar excess of succinimidyl 4-[N-maleimidomethyl]cyclo-hexane-1-carboxylate (SMCC-Sigma-Aldrich #M5525) for 2.5 hours at room temperature to generate maleimide-DNA capable of sulfhydryl-reactive crosslinker chemistry with peptides with terminal cysteines. SMCC-Oligo conjugates were isolated using ethanol precipitation and dissolved in T150 buffer (25 mM Tris, 150 mM NaCl, 1 mM EDTA), and then run through Bio-Spin 6 columns (Bio-Rad #7326002) twice. Peptides with terminal cysteines were synthesized by GenScript and diluted to 200 µM in T150 buffer: WT (C- NANDPAPPNANDPAPPNANDPAPP-C) or Scrambled (C-NNNAPDDNAAANPPANPPPPPD-C).

Peptide was added to SMCC-Oligo at a 1:2.5 molar ratio and incubated overnight at 4°C for conjugation. Unreacted SMCC-Oligo was removed by polyacrylamide gel-electrophoresis followed by extraction from the gel. 250 pmoles of oligo-peptide conjugate was added to 250 pmoles 5’ -biotin- TGGCGACGGCAGCGAGGC-Cy5- 3’ and 300 pmoles 5’ –GGGCGGCGACCTGCTGGGTAGTC-Cy3- 3’ and incubated overnight at 4°C for conjugation by complementary base pairing with the DNA conjugated to the peptide. For fluorescence force experiments, 16 nM λ-DNA (NEB #N3011S) in 0.120 M NaCl was heated to 80°C for 10 minutes followed by 5 minutes on ice. The Cy3-Cy5- peptide construct was added at 10 nM with 0.2 mg/mL BSA and incubated at 4°C for 3 hours followed by addition of 200 nM 5’ -AGGTCGCCGCCCTTT- digoxygenin- 3’ and overnight incubation at 4°C. For single-molecule fluorescence-force spectroscopy experiments, an imaging chamber was made by sandwiching polyethylene-glycol (PEG) (a mixture of PEG-valeric acid and biotin-PEG-valeric acid, Laysan Bio) between a passivated quartz slide and coverslip. The imaging chamber was then incubated with 0.2 mg/mL neutravidin (Pierce) in blocking buffer (10 mM Tris-HCl pH 7.5, 50 mM NaCl, 1 mg/mL BSA, 1 mg/mL tRNA (Ambion)) for 1 hour. The peptide constructs were then diluted to 20 pM and immobilized on the surface via biotin-neutravidin interaction. 1 M anti-digoxygenin coated polystyrene beads (Polysciences) were conjugated to the peptide construct through λ-DNA of the peptide construct by incubation in a buffer containing 10 mM Tris HCl pH 7.5 and 150 mM NaCl for 30 minutes. Microscope setup consisted of a trapping laser (1064 nm, 800 mW, Spectra Physics) to catch a bead. As the microscope stage was moved along the X- plane, a confocal laser (532 nm, 30 mW at maximum average power (World StarTech)) was used to follow Cy3/Cy5 emission profiles. To apply tension, the stage was moved between 14 and 16.8 µm at a constant rate of 445 nm/sec as Cy3/Cy5 emission profiles were measured with an exposure time of 20 milliseconds. The applied force was measured by a quadrant photodiode (UDT/SPOT/9DMI) from the position of the tethered bead. FRET trajectories as a function of force applied to peptides were then binned by 0.5 pN increments and plotted using Origin software (OriginLab).

### Statistical Analysis

Statistical analysis were performed in GraphPad Prism. All data were evaluated for skewedness using Graph Pad Prism’s statistical analysis. Datasets with skewedness values <-1 or >1 were considered non-parametric. For comparison between two groups either Mann-Whitney tests (non-parametric data) or Welch’s t-tests (parametric data) were performed. For comparisons between multiple groups, a one-way ANOVA followed by Dunnett’s post hoc test (parametric data) or Kuskal-Wallis followed by Mann-Whitney-Wilcoxon (non-parametric data) was used. For proportion data, a Z-score for population proportions was performed using the equation 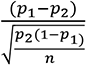 with significance at p<0.01. The tests used and p-values are described in the figure legends.

## Acknowledgements

We are grateful to the Insectary and Parasitology Core Facilities at the Johns Hopkins Malaria Research Institute, expertly staffed by Chris Kizito, Godfree Mlambo and Abai Tripathi, and to the Bloomberg Family Foundation for their support of this facility. We are grateful to Barbara Smith and Dr. Scott Kuo of the Johns Hopkins University School of Medicine Microscope Facility for their invaluable assistance with the RICM imaging experiments. This study was supported by a fellowship from the Johns Hopkins Malaria Research Institute (AB), the National Institutes of Health grants R01AI056840 (PS) and R35 122569 (TH), and the Bloomberg Family Philanthropies. The RICM imaging reported in this publication was supported by Office of the Director of the National Institutes of Health under award number S10OD016374.

---

**Supplementary Table 1.**
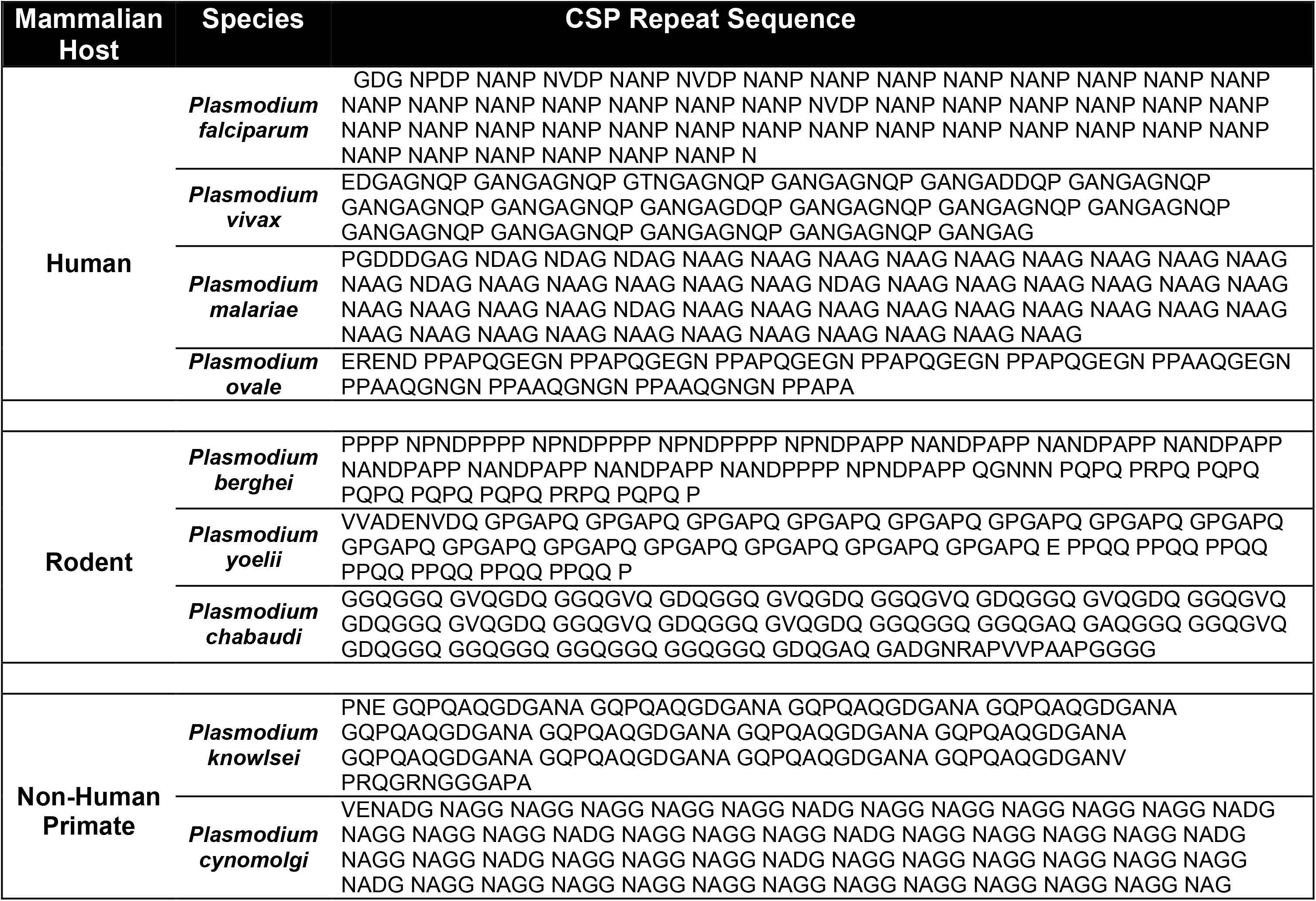
Comparison of CSP Repeat Sequences Across *Plasmodium* Species. Within each sequence the individual repeat blocks are separated for illustrative purposes. We define the repeat region as beginning just after the highly conserved Region I: In most cases there is an initial 3 to 9 amino acid non-repetitive sequence as shown.

**Supplementary Figure 1.**
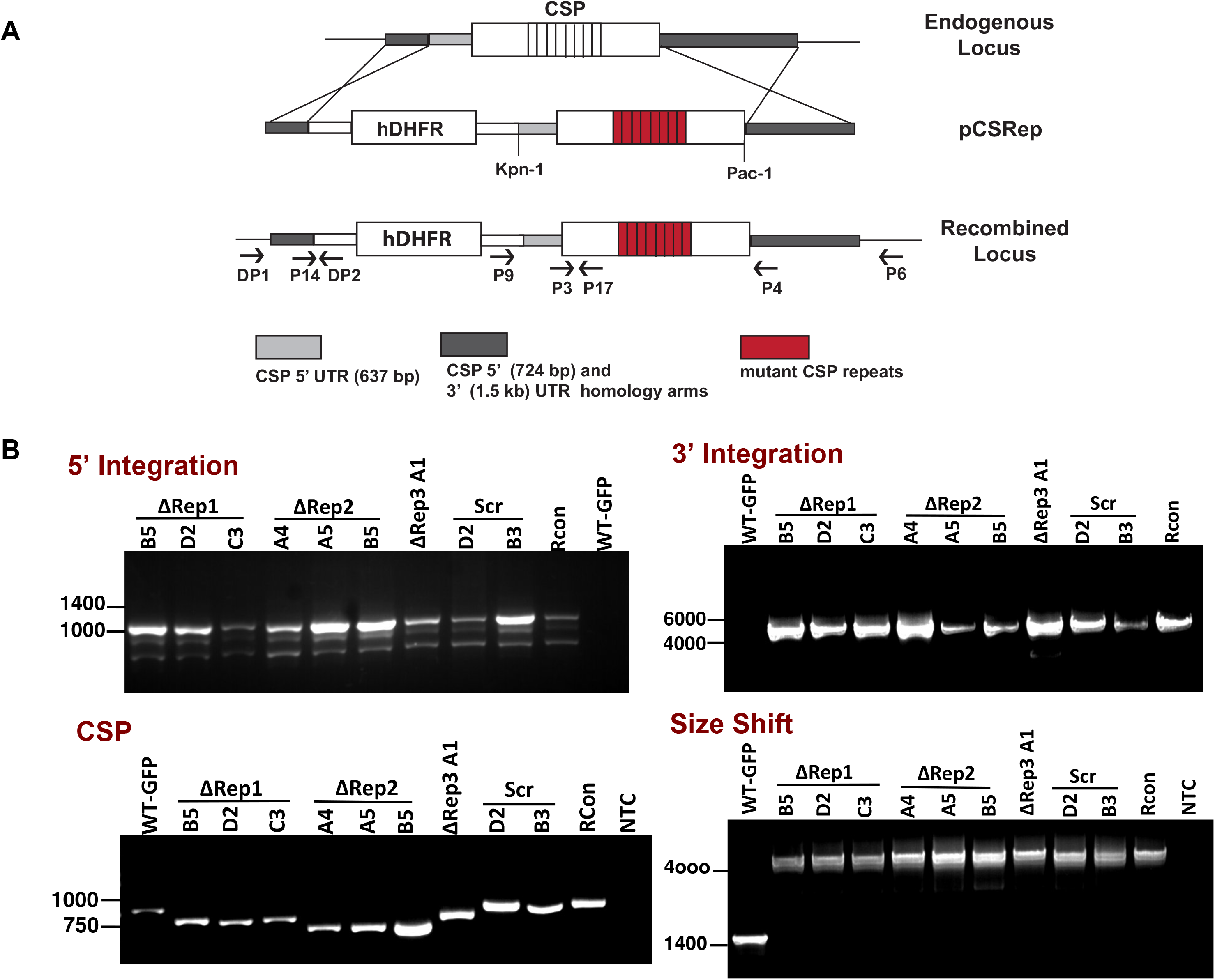
Generation and validation of CSP repeat mutants. (**A**) Transfections were performed to replace the endogenous *csp* locus with a *csp* gene that contained a repeat region with the desired mutations. The indicated fragment from pCSRep was released from the plasmid by restriction digest, with the 5’ and 3’ UTR homology arms driving homologous recombination to give rise to the recombinant locus shown. The hDHFR selection cassette upstream of *csp* enabled positive selection of recombinant parasites. Binding sites of primers described in the genotyping panels below are also shown. (**B**) Clonal lines were verified using a series of PCR reactions: DP1 and DP2 for 5’ integration, P9 and P6 for 3’ integration; P3 and P4 for the *csp* gene with size changes due to truncation and P14 and P17 which shows a size shift in recombinant parasites due to the insertion of the hDHFR cassette.

**Supplementary Figure 2.**
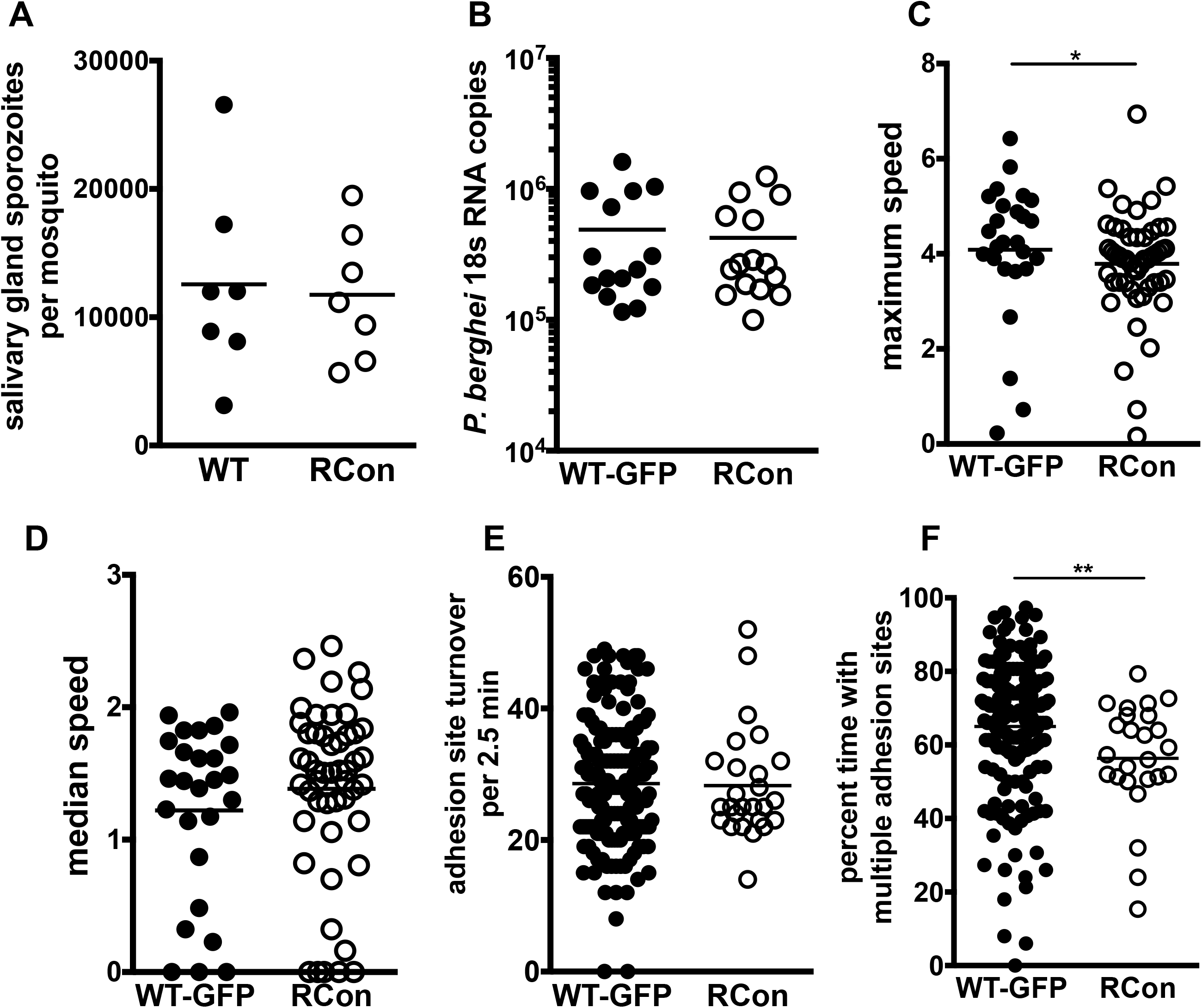
Recombinant control (RCon) and WT-GFP sporozoites have similar phenotypes. (**A**) Salivary gland sporozoite loads. Salivary glands from 20 infected mosquitoes were pooled, homogenized and the average number of sporozoites per mosquito was determined. Each data point is from an independent mosquito cycle with matched WT-GFP and RCon lines and data are pooled from 7 biological replicates. Paired Wilcoxon t-test at p<0.05 showed no statistically significant difference. Welch’s t-test p = 0.8168. (**B**) Sporozoite liver infection. 5,000 WT-GFP or RCon sporozoites were inoculated IV into C57Bl/6 mice and 40 hours later, parasite liver burden was determined by RT-qPCR using primers specific *P. b*erghei 18s RNA. Shown are pooled data from 3 experiments with 5 mice per group in each experiment. (**C&D)** Live gliding assay showing maximum (C) and median (D) sporozoite speeds from 26 (WT-GFP) and 50 (RCon) sporozoites. For (C): Mann-Whitney-Wilcoxon *p=0.047. For (D) : Welch’s t-test p = 0.3036. (**E**&**F**) Adhesion site analysis using RICM. RICM movies of circular gliding WT-GFP and RCon sporozoites were analyzed to quantify the frequency with which adhesion sites move from the sporozoite’s anterior to posterior end (E) and the frequency with which gliding sporozoites have multiple adhesion sites (F), Mann-Whitney-Wilcoxon **p = 0.004. All movies included 2.5 minutes of circular gliding for analysis. Data are pooled from at least 3 independent experiments with the following number of sporozoites per line: WT-GFP=175 and RCon = 25.

**Supplementary Figure 3.**
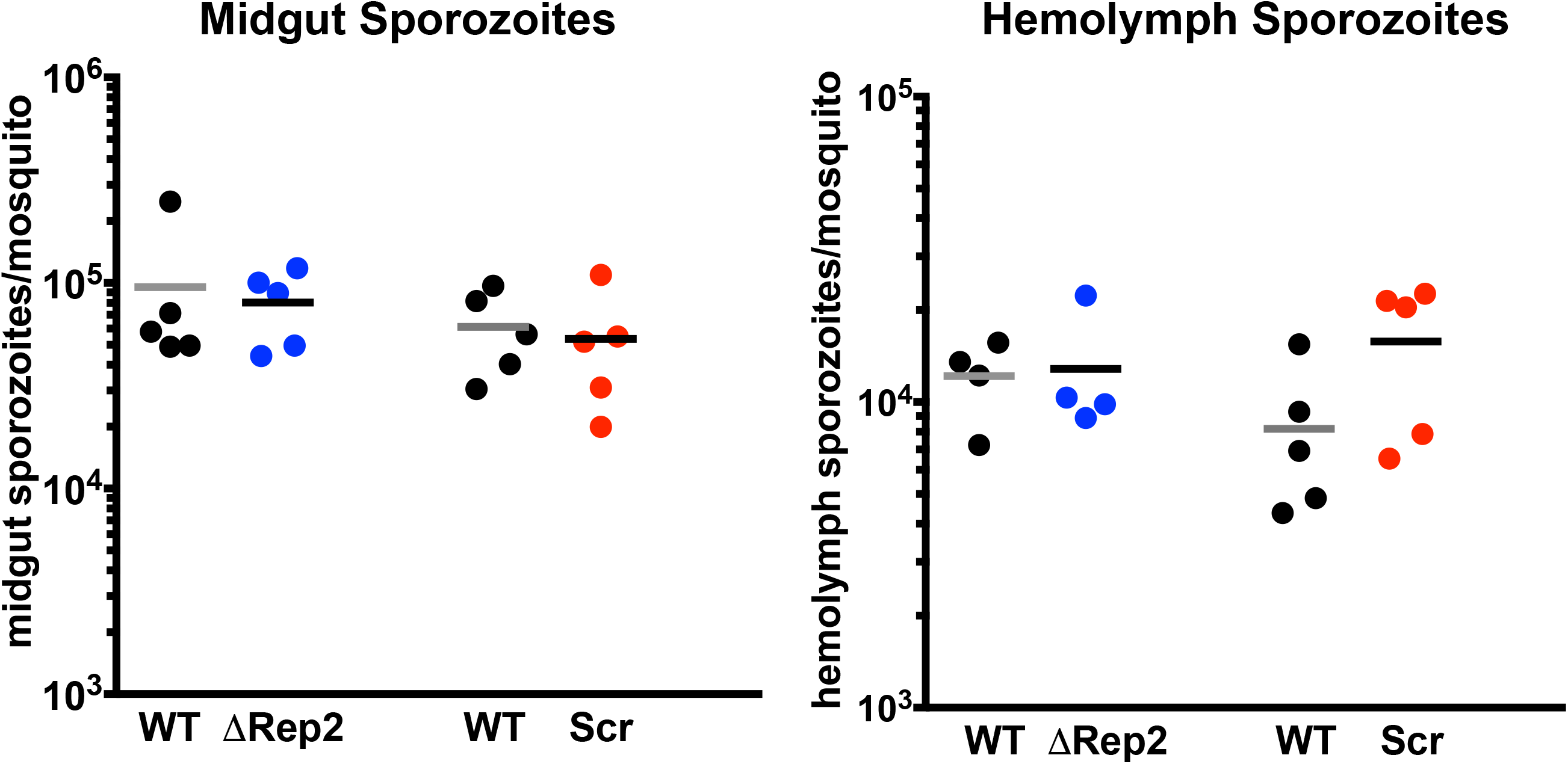
CSP Repeat mutants exhibit normal sporozoite development in the mosquito prior to salivary gland invasion. Midgut (left panel) and hemolymph (right panel) sporozoites were collected from 20 mosquitoes and counted. Shown is the average number of midgut or hemolymph sporozoites per mosquito. Each data point is from an independent mosquito cycle with matched control lines and shown are pooled data from 5 to 6 biological replicates. Kruskal-Wallis showed no significant differences between mutant lines and matched controls (midgut sporozoites, p=0.6026; hemolymph sporozoites, p=0.2979).

**Supplementary Figure 4.**
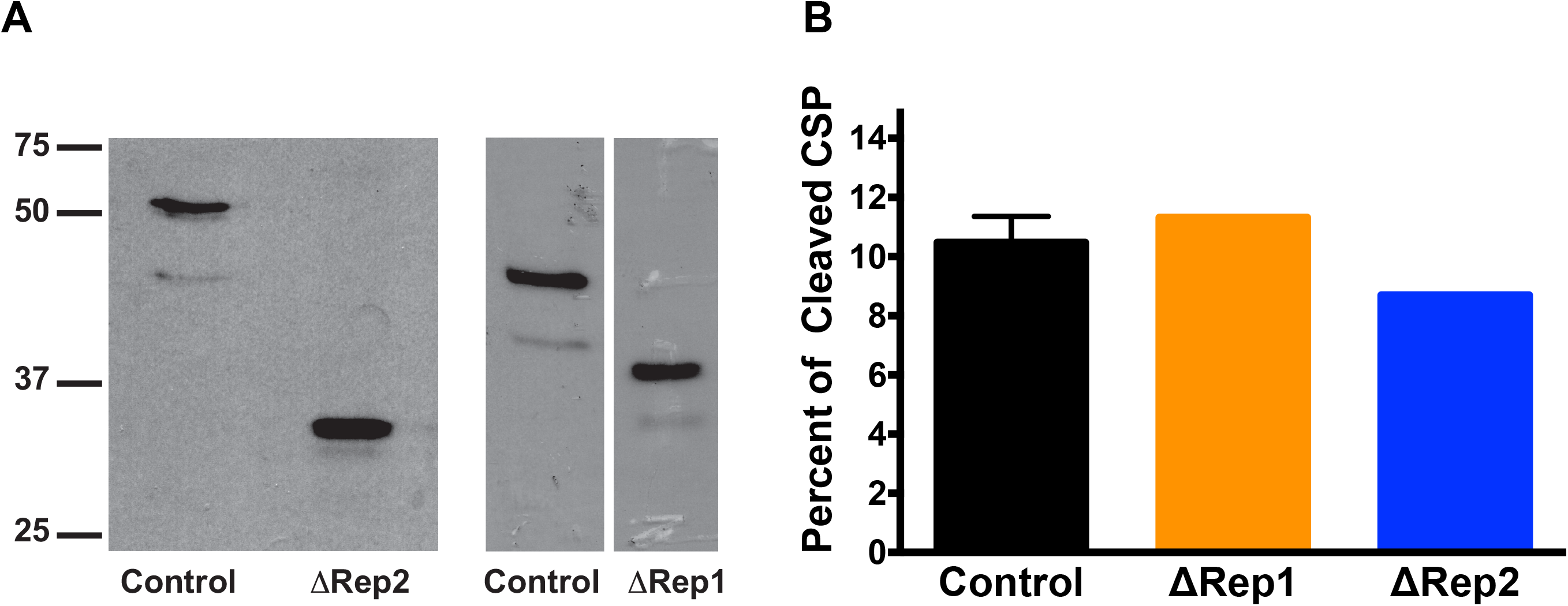
TSR exposure in. Δ**Rep2 is not due to enhanced CSP cleavage. (A**) Control, ΔRep2, and ΔRep1 sporozoites were metabolically labeled with [35S]-Cys/Met and CSP was immunoprecipitated and run on an SDS-PAGE gel followed by autoradiography. (**B**) Band intensities were measured by densitometry and percent of total CSP detected in the bottom band (cleaved) is expressed in the graph in panel B. Densitometry measures of control parasites are pooled from both blots.

**Supplementary Figure 5.**
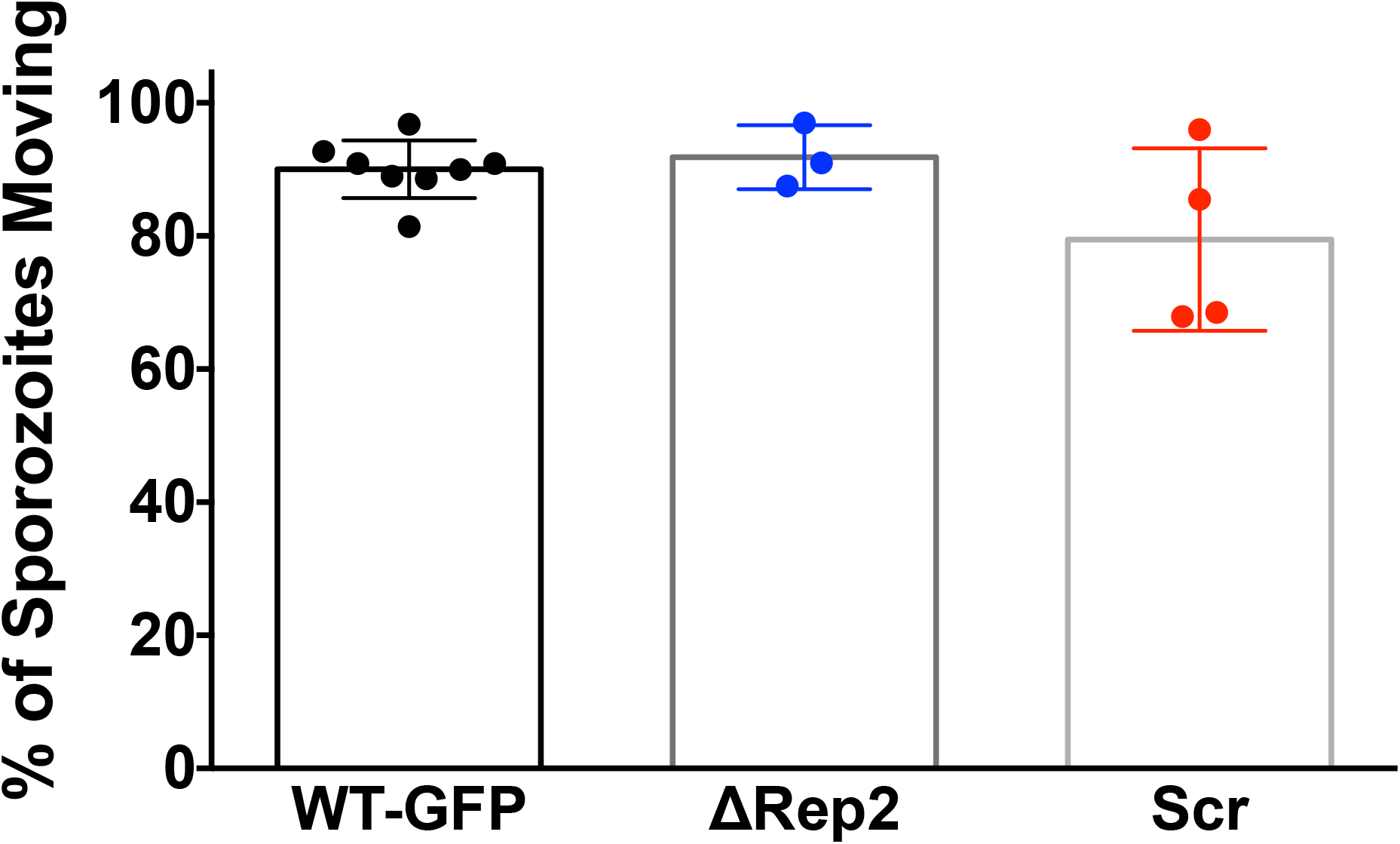
The percentage of CSP repeat mutants that are motile is similar to controls. Sporozoites were prepared for live gliding experiments, added to coverslips and at least 50 sporozoites per coverslip were counted. Each sporozoite was recorded as moving (circular gliding, patch gliding or attached waving) or not moving (drifting or attached with no movement). Any attached sporozoite that was not moving was observed for at least 30 seconds prior to being recorded as non-motile. Data are pooled from 3 to 4 biological replicates with each dot representing the percentage of sporozoites moving from one experiment with pooled controls that were performed alongside mutants. Comparison of percent motile between the control and ΔRep2 or Scr line was not significant; Kruskal-Wallis test p=0.2649.

**Supplementary Figure 6.**
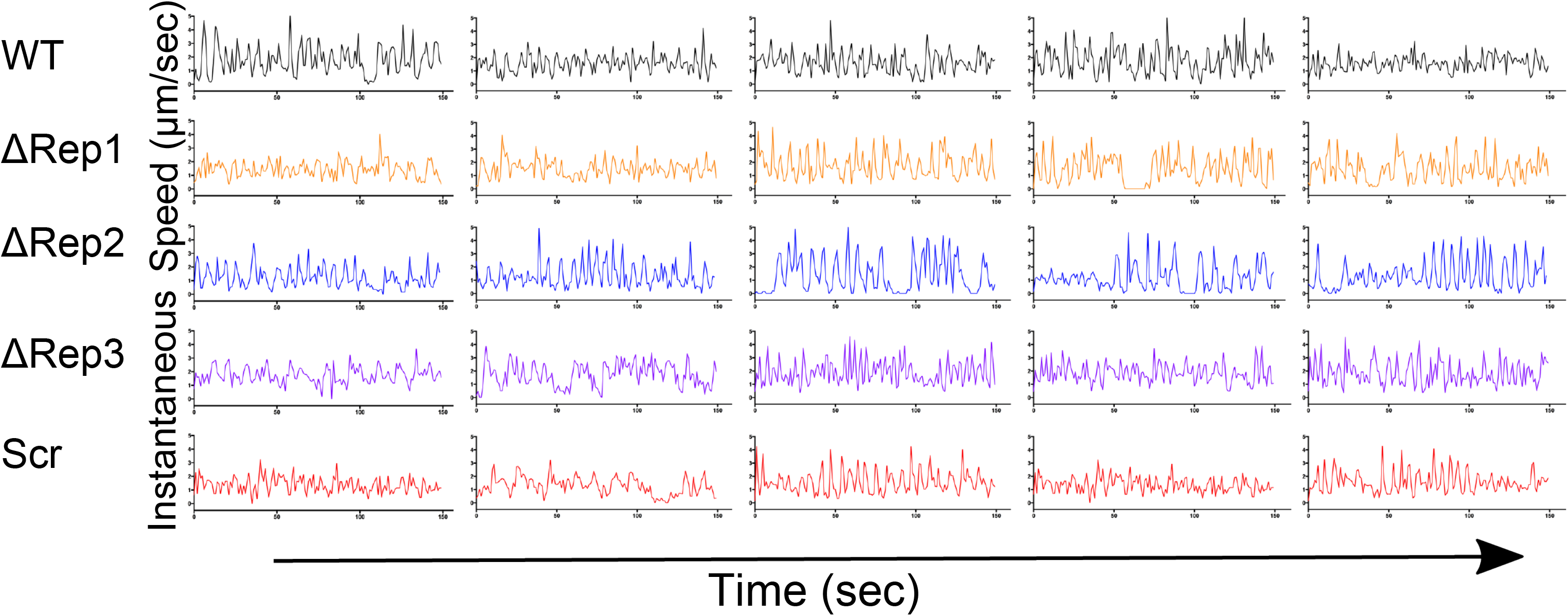
Instantaneous speed plots of circular gliding CSP repeat mutants. Five representative instantaneous speed plots are shown. Plots were chosen such that each had a median speed close to the mutant’s overall median speed (shown in Figure 5B). Both x- and y-axes are identical for all plots.

